# The RIFF: an automated environment for studying the neural basis of auditory-guided complex behavior

**DOI:** 10.1101/2021.05.25.445564

**Authors:** Maciej M. Jankowski, Ana Polterovich, Alex Kazakov, Johannes Niediek, Israel Nelken

**Author notes:** These authors had equal contributions (Maciej M. Jankowski, Ana Polterovich, Alex Kazakov, Johannes Niediek).

## Abstract

Behavior consists of the interaction between an organism and its environment, and is controlled by the brain. Brain activity varies at sub-second time scales, but behavioral measures are usually coarse (often consisting of only binary trial outcomes). To overcome this mismatch, we developed the RIFF: a programmable interactive arena for freely-moving rats with multiple feeding areas, multiple sound sources, high-resolution behavioral tracking, and simultaneous electrophysiological recordings. We describe two complex tasks implemented in the RIFF. Rats quickly learned these tasks and developed anticipatory behavior. Neurons in auditory cortex and posterior insula showed sensitivity to non-auditory parameters such as location and pose. Our combination of wireless electrophysiology and detailed behavioral documentation in a controlled environment produces insights into the cognitive capabilities and learning mechanisms of rats and opens the way to a better understanding of how brains control behavior.

## Introduction

Behavior involves brain-controlled, complex interactions between an organism and its environment. Natural behavior has many components. Much of what we understand about brains and their functions was shaped by examinations of single components of the interaction between organisms and their environments, including sensation and perception (Schnupp et al., 2010); motor control (Shadmehr et al., 2010); decision making (Carandini & Churchland, 2013); and memory (Poo et al., 2016). However, these successes have been achieved at a price – animals are often studied using behavioral tasks that are much simpler than those they face in their natural habitats.

The main observable output of behavior is movement – animals select and time their actions. Patterns of movements may be rich: predictive movements may precede trial events when there is a temporal structure to the task; movements may differ between individuals; and movements may be used for solving cognitive tasks. For example, rats exert active control on selected motor variables in order to enable consistent perception of location when using their whiskers (Saraf-Sinik et al., 2015).

Evidently, even in simple tasks, animals perform much more than a single action at each trial. In complex, naturalistic settings, the amount of movement (and number of decisions) performed by animals is substantial. Brain activity is expected to reflect and shape the full complexity of the concomitant behavioral strategies (Musall et al., 2019).

Nevertheless, brain activity and behavior are often measured at very disparate rates. While brain activity changes at rates reaching 100/s or faster, behavior is often quantified by coarse measures at the rate at which trials are presented. Response latencies and outcome-driven measures do not lend access to the entire decision process occurring from trial onset to outcome. This is a substantial limitation. During a trial, neural activity throughout the brain runs its fast course, before, during and after relevant decision points. For example, movements that vary from trial to trial account for much of the inter-trial variability in wide-field calcium imaging of mouse cortex (Musall et al., 2019). Thus, a large amount of information in the neural responses about behavior, which holds valuable insights on the links between behavior and neuronal processing, is likely wasted.

We designed and constructed the Rat Interactive Foraging Facility (RIFF) that allows high-resolution behavioral data acquisition in time and space combined with tetherless electrophysiology in freely moving animals. The RIFF is controlled by a real-time loop that tracks the rat, identifies its actions and reacts to them. Because of the generic nature of this loop, it is flexible and can implement a rich set of behavioral scenarios.

We describe here the design principles of the RIFF and their implementation using two very different tasks. Rats learned these behavioral tasks efficiently, sometimes within a few hours. Different rats developed individual patterns of behavior, which can be detected early in the learning process and tracked over months. We show that neural activity recorded in primary and higher-order auditory fields tracks relevant behavioral parameters such as animal location and pose in addition to responding to auditory stimuli.

## Results

### Overview of the RIFF

The RIFF is designed to jointly study behavior and brain activity of freely-moving animals interacting with a rich environment that is nevertheless amenable to good experimental control. The implementation of the RIFF requires integrating a large number of interacting systems and processes (Fig. 1), which we kept as separate and modular as possible in order to allow for modifications and extensions as experience is gained and new technology becomes available. While the logic used to operate the RIFF can be very flexible, we intended it to operate mostly as a Markov Decision Process (Sutton & Barto, n.d.) (MDP). MDPs are ‘state machines’. They are defined by a set of states (defined, for example, by the location of the animal, the current stimulus that is presented, and the correct ports to poke in order to get a reward), a set of actions that an animal can take in each state, and a set of actions that can be taken by the environment (typically providing rewards and punishments to the animal). The MDP is governed by two (potentially stochastic) rules. The first is the rule by which one state follows another. In an MDP, transitions depend only on the current state of the process and on the current action of the animal. The second rule prescribes whether and which actions the environment takes, depending only on animal action and the consequent state transition.

**Fig. 1.**
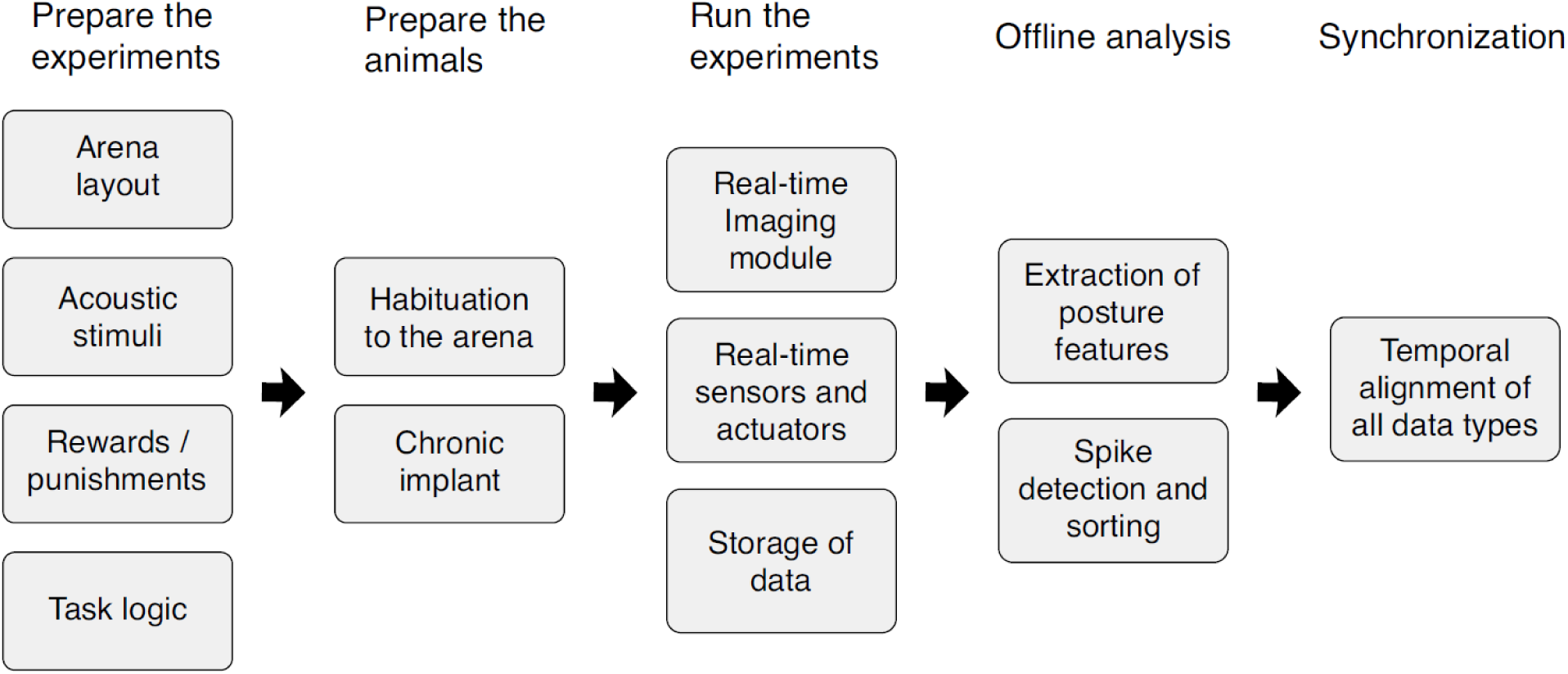
Workflow for the design, preparation, execution and initial analysis of behavioral experiments as they pertain to the RIFF. The design of an auditory-guided task includes the development of task logic, the nature of rewards and punishments, as well as the selection of the auditory stimuli. The flexibility of the RIFF allows for substantial freedom in these choices. Rat preparation includes habituation to the RIFF and electrode implantation. While running the experiment, the RIFF is designed to keep a tight synchronization of all real-time events, and to ensure efficient data storage. The stored data allows for offline analysis that includes spike sorting and animal pose estimation, which are then precisely aligned with the rest of the information from the experiment.

The behavioral environment is a large 18-sided polygon (Fig. 2a) with six interaction areas (IAs), each consisting of two loudspeakers, a water port, a food port, and air-puff valves (Fig. 2b). The main sensor for animal location is a ceiling-mounted camera with a dedicated computer that performs real-time image processing. The camera tracks the location of the rat with a temporal resolution of 30 ms, and the location information is transmitted to the main computer. Additional sensors include the nose-poke detectors at each feeding port that report their state in real time; the status of the food and water ports; and analog sensors carried by the animals such as a microphone, accelerometers, and gyroscopes (Mallory et al., 2021). The RIFF interacts with the behaving animal through the actuators (Fig. 2b). In the present implementation, the response repertoire of the RIFF includes food or water delivery, air-puff delivery, and sound presentation by any combination of the 12 speakers. Neural responses are recorded using chronically implanted silicon probes (Fig. 2c).

**Fig. 2.**
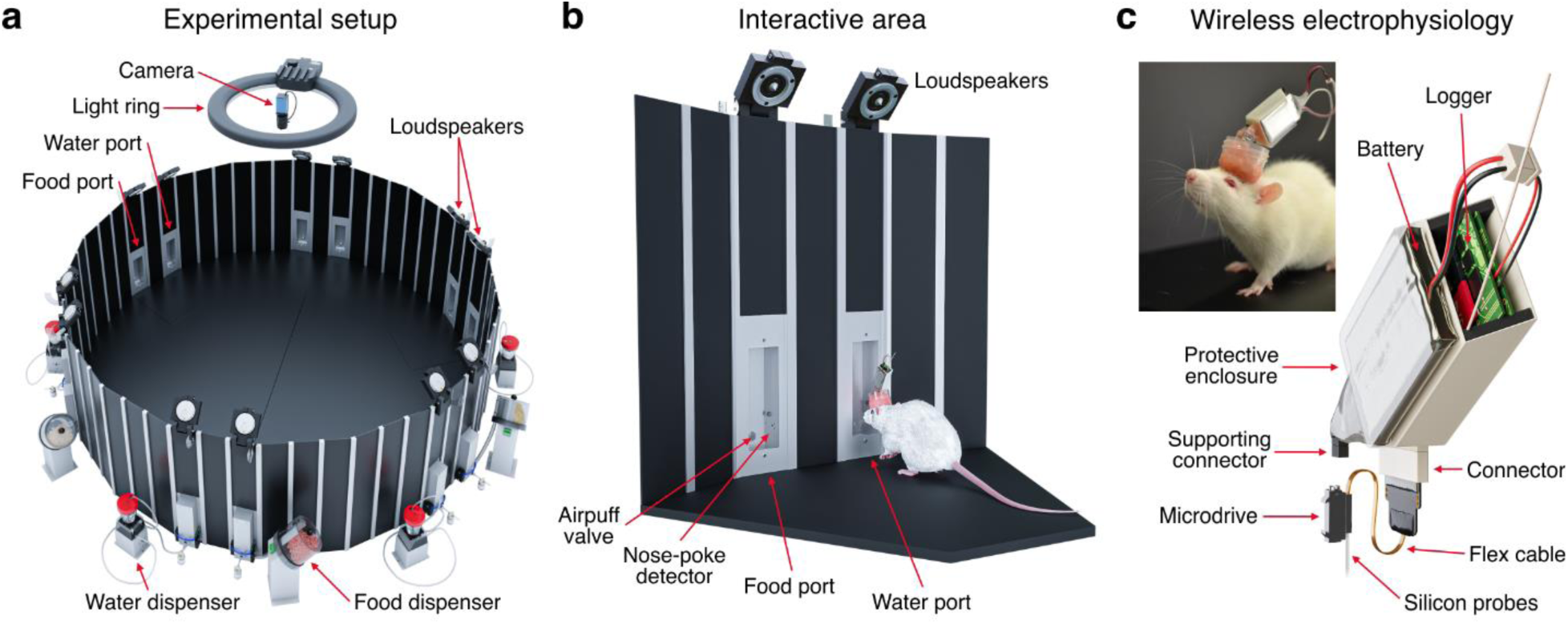
(a) An experimental arena for studying the neural basis of auditory-guided complex behavior. The RIFF is a large arena (160 cm in diameter), equipped with 12 loudspeakers, 6 food dispensers and 6 fluid dispensers, 12 airpuff outlets, and a ceiling mount camera. The wall panels can be removed, the ports can be closed by blinds, and the spatial arrangement of the environment can be adjusted for different experiments. **(b)** Close-up on one interaction area. An adult female rat with a chronic implant and a neural logger is shown for scale. Each interaction area consists of two ports, one for food rewards and the other for fluids. Each port has a nose poke detector and airpuff outlet. Two speakers are mounted above each port. **(c)** An approach for chronic wireless recordings in rats. The neural logger and battery are in the protective plastic case. A multi-contact silicon probe is mounted on the microdrive. The device is inclined to the back in order to allow rats to move naturally and an undisturbed access to the ports of the interaction area. The picture shows a rat with a 32-channel moveable silicon probe implant and wireless data logger. The battery used here allows for 3 hours of recordings.

The logic of the RIFF is implemented in a program that runs on the main control computer (Fig. 3a), collecting the data from sensing hardware, and driving the actuators using a predefined logic (an example is described in Fig. 3c; see also Methods and Supp. Fig. S1). Four major data types are recorded during the experiment and stored for offline analysis (Fig. 3b): time stamps, neural activity, analog sensor data, and images.

**Fig. 3.**
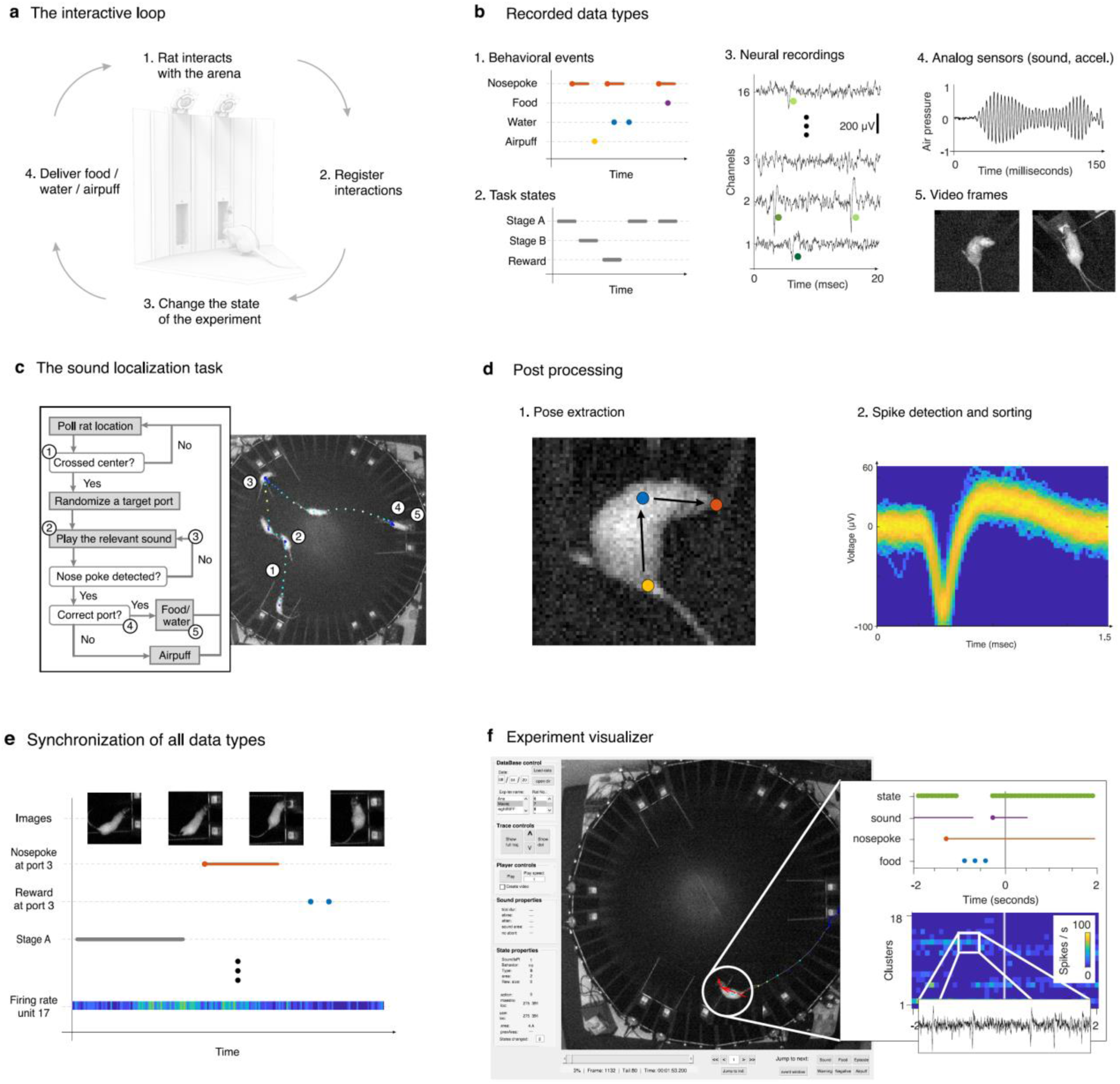
Operation of the RIFF. **(a)** Rats interact with the RIFF through movement and nose pokes. Their location is identified online using the video camera and a dedicated computer. The experimental environment reacts to rat actions by changing its state and providing different types of feedback – sounds, rewards, or punishments. **(b)** The RIFF collects multiple data types: behavioral events (e.g. nose pokes, rewards, punishments); task states; neuronal activity; analog sensor signals (microphone, motion sensors); and the animal location tracked by the ceiling mount camera. **(c)** An illustration of a sequence of interactions between a rat and the RIFF during the LD task. (1) The rat moved to the center of the arena in order to initialize the trial. (2) Once it crossed into the central area (a circle with a radius of 30 cm), the RIFF started sound presentation from both loudspeakers in a randomly selected interaction area. (3) Initially, in the example presented here, the rat approached a wrong port; a second sound presentation caused it to move towards the correct target port (4), and to receive the reward (5). **(d)** Data post-processing. Posture features are extracted using a custom-trained DNN. The Kilosort2 program used with a custom wrapper performs largely automatic spike detection and sorting. **(e)** All data types are synchronized on a single time axis and **(f)** can be visualized offline using a custom visualizer software. The visualizer can be used to browse through all synchronized data types up to the level of the raw neural signals.

The RIFF is designed for high-throughput data acquisition with almost completely automated post-processing steps. A convolutional neural network estimates the locations of the nose, neck and the base of the tail of the rat from the video stream, inferring head and body directions (Fig. 3d.1, Supp. Fig. S2, and Supp Movies S1 and S2). The neural recordings are denoised, and spike detection and sorting is performed by a custom wrapper to Kilosort2 (https://github.com/MouseLand/Kilosort2) (Fig. 3d.2, Supp Fig. S3). After extracting information from each data stream, all data are synchronized to a single time axis (Fig. 3e). The output is a single file that combines all data types and additional extracted features. The post-processing steps are much faster than actual experiment time. An interactive graphical interface (Fig. 3f, Supp Movie S3), is used to verify the experiment logic and data integrity, and to examine the fine details of the behavior and the data.

### Rats rapidly learned a complex task

To illustrate the flexibility of the RIFF, we implemented two different behavioral tasks: a multiple-strategy (St+) task and a localization/discrimination (L/D) task (Both tasks are described in detail in the Methods section; see Supp. Fig. S1 and Supp. Movies S4 and S5). Both tasks were run daily in the same setup on different subjects.

We illustrate rapid learning in the RIFF using the St+ task. Each behavioral episode of the St+ task started with an attention sound, followed by a period of 2.5 seconds during which the rat could control the selection of the IA for the next reward by moving to appropriate positions in the arena. Next, this reward location was communicated to the rat by a target sound emitted by the loudspeakers of the selected IA. The rat had to poke in one of the two ports of the target IA within 20 s of the presentation of the target sound. A poke or a timeout resulted in a feedback sound followed by an inter-trial interval (3 s) until the next attention event. Rats rapidly learned the task despite its non-trivial structure (see Supp. Movie S4). We analyze here in detail the first two days of task exposure.

Each rat trained in the RIFF for 70 minutes per day. During the first two days of training, rats quickly learned to move from one IA to the next. Figure 4a shows one-minute segments of a rat’s trajectory on Day 1 and Day 2. Erratic loops and hesitations were present on Day 1 but disappeared on Day 2. Figure 4b shows the angular position and angular speed from the same trajectories, along with a schematic illustration of the trial structure. Clearly, the stereotypic running pattern of Day 2 was still absent at the beginning of Day 1.

**Fig 4.**
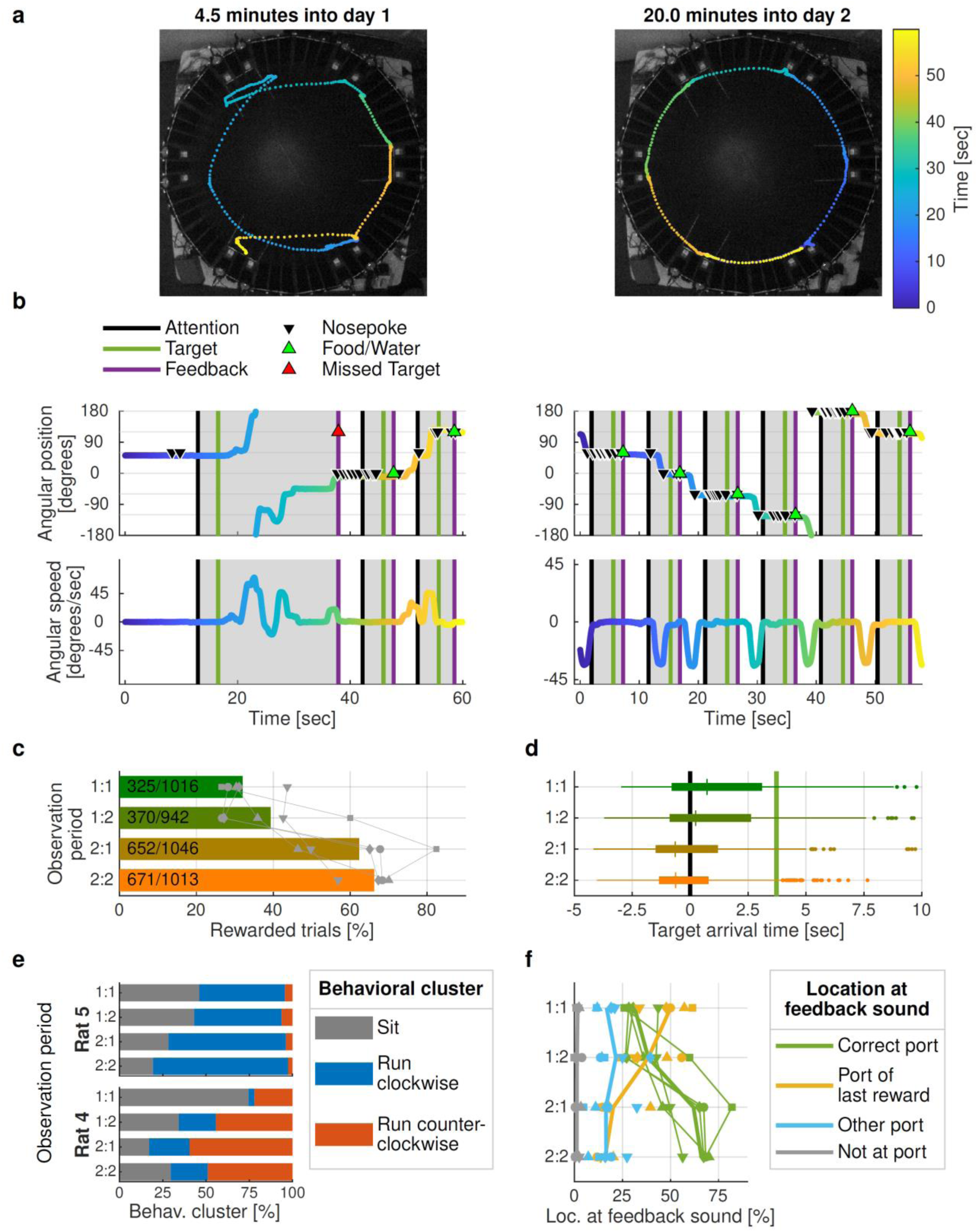
Rapid learning in the RIFF. **(a)** One-minute trajectory segments from 70-minute training sessions on days 1 and 2 of learning. **(b)** Angular position and speed for the same trajectories. Trials (from attention to feedback) are marked in gray. Nosepokes, rewards, and missed targets are indicated by triangles. By day 2, rats have developed a stereotypic running pattern. **(c)** Success rate improved over time. Observation periods 1:1/1:2 denote the first/second half of the session on day 1; 2:1/2:2 denote session halves on day 2. Bars indicate average success rates from five rats; gray symbols mark individual rats. **(d)** Rats learned the temporal structure of the task. Shown are arrival times relative to the attention sound (black line; target sound, green line), calculated from all trials of five rats. **(e)** Different rats learned different strategies. Each trial was classified as either sitting, clockwise running, or counterclockwise running. Rat 4 preferred counterclockwise running, while Rat 5 avoided counterclockwise running. **(f)** Learning to move was the most important contribution to performance improvement. Each trial was classified as either “correct port”, “port of last reward”, “other port”, or “not at port”. Average proportions of each location across five rats are displayed as thick lines, individual rats are displayed as symbols and thin lines. The proportion of “correct” locations increased from one observation period to the next, and the proportion of “port of last reward” locations decreased over time.

To analyze changes in behavior within and across sessions, we subdivided each session into two halves, denoting the four resulting observation periods by 1:1, 1:2, 2:1, and 2:2. The percentage of rewarded trials - a simple measure of learning - increased from 32% in observation period 1:1 to 66% in observation period 2:2 (Fig. 4c). We modeled the percentage of rewarded trials as a function of the observation period using a linear mixed-effects (LME) model with random intercepts for rats. The model had a significant main effect of observation period with a slope corresponding to an average improvement of 13% ± 2% (mean ± STE) additional rewarded trials from one observation period to the next (t(18) = 5.95; p = 1.2*10^-5^). As rats learned to position themselves to collect rewards, they adapted to the timing of the task: rats arrived in the IA of an upcoming reward one second earlier on Day 2 compared to Day 1 (Fig. 4d; median arrival time for successful trials on Day 1, 0.4 seconds (interquartile range (IQR) 3.7 s) after attention sound; on Day 2, 0.6 seconds (IQR 2.4 s) before attention sound).

This suggests that rats learned to correctly predict the upcoming reward time. We modeled the logarithm of arrival times as an LME with the observation period as a fixed factor and with random intercepts for rats (times were shifted to start 5.5 s before the attention sound to turn all arrival times into positive numbers; the logarithmic transformation was selected in order to reduce the skewness of the distribution of arrival times). There was a significant effect of observation period (estimate -0.069 ± 0.008; t(1829) = -8.86; p = 1.8*10^-18^), so that the arrival time shifted to earlier times by a factor of exp(-0.069 ± 0.008) = 0.93 ± 0.007 from one observation period to the next.

The velocity trajectories in Fig. 4a suggest that the movement patterns of the rats quickly became stereotypical. We calculated the maximal angular speed in both the clockwise and counterclockwise directions during each trial (analysis time window, 0.5 to 9 seconds after the feedback sound of the previous trial). All rats learned three clearly distinct behavioral motives: “not moving”, “clockwise running”, and “counterclockwise running” (Supp. Fig. S4 for cluster analysis). Each experimental session can thus be described as a sequence of these three motives, timed by the rat to harvest rewards from the RIFF.

We analyzed how the distribution of the three motives evolved during the four observation periods. There was a large decrease in the number of trials in which the rat did not move. We modeled the proportion of “not moving” trials as a linear function of the observation period using an LME with random intercepts for rats. Observation period had a significant main effect (estimate, -0.14 ± 0.02; t(18) = -5.89; p = 1.4*10^-5^). Thus, on average, the proportion of “not moving” trials decreased by 14% ±2% from one observation period to the next: rats learned to move more often as training progressed.

Importantly, rats showed a clear, individual preference for running in one direction or the other. Thus, we found that the proportion of “clockwise running” out of all running trials depended on the rat (two rats shown in Fig 4e; χ^2^-test for independence; χ^2^ (24) > 701 for the five rats together; p < 10^-20^). In both rats, the fraction of “not moving” trials (shown in gray in Fig. 4e) generally decreased over time. Rat 4 preferentially ran counterclockwise (blue), while rat 5 preferentially ran clockwise (red). The distribution of “clockwise”/”counterclockwise” significantly depended on rat identity (these two rats: χ^2^(21) > 77.6, p < 10^-20^ in each observation period).

Lastly, to better understand the nature of failed trials, we classified the rat location at the time of the feedback sound into four types: “correct port” (rewarded trials), “port of last reward”, “another port”, and “not at any port”. As training progressed, rats reached the correct port more often (green line in Fig. 4f). The most common error type consisted of remaining in a reward location after a reward, thereby missing the next reward opportunity which was always at a different port (orange line in Fig. 4f). This type of error decreased substantially from Day 1 to Day 2. Statistical analysis confirmed that the distribution of rat location types depended on the observation period (χ^2^ (245) = 617, p < 10^-20^ for all rats; χ^2^ (29) > 38.4, p < 1.5*10^-5^ in each individual rat). To quantify the interaction between the location types and observation periods, we modeled the probability of the location types as an LME model in which the probability for each location type was a linear function of the observation period with a slope that depended on the location type, with random intercepts for each location type in each rat. This model revealed a significant interaction between observation period and location type (F(3,72) = 44.1; p = 2.8*10^-16^), confirming that over time, the rats changed their preferred location types. Indeed, the slope of the “correct port” location type was 0.13 ± 0.02 (t(72) = 8.36; p = 3.2*10^-12^), confirming the increase in correct trials over time. In contrast, the slope for “port of last reward” location type was -0.12 ± 0.02, showing a decrease in this type of error. The difference between these two slopes was highly significant (t(72) = -11.5; p = 6.1*10^-18^).

### Neural correlates of behaviors

Rats in the St+ task were implanted with electrodes targeting the auditory field in the posterior insular cortex (Ins), which in rats is anatomically separate from other auditory cortical fields (Kimura et al., 2010; Rodgers et al., 2008). Rats in the L/D task were implanted with electrodes which traversed the primary auditory field at least in part of their trajectory (AC).

Units in both the AC and Ins showed responses to sounds. Figures 5a and 5b show responses of units in AC of two rats. The stimuli were word-like stimuli (200 ms word excerpts, with their spectral envelopes shifted and stretched to fit a 1-40 kHz range; see Methods). These stimuli were some of the target sounds in the L/D task (mean response of N = 96 and N = 143 sound presentations in Figs 5a and 5b respectively; here and elsewhere in this paragraph a paired t-test was performed between spike counts in the response window and in a window of the same duration just preceding stimulus onset, t(190)=3.26, p = 0.0013 and t(284)=2.63, p = 0.0090 respectively). Both units had well-timed onset responses. Figures 5c and 5d show responses of units in Ins. The stimulus in Fig. 5c was the attention sound (a train of short broadband periodic sound bursts with a pitch of 2 kHz). The unit responded to the onset of the first sound burst in the train (mean of N = 233 sound presentations, t(464) = 4.48, p = 9.6*10^-6^). Figure 5d shows a unit with a response that was loosely locked to a word-like stimulus (mean of N = 29 sound presentations, t(56) = 3.27, p = 0.0018).

**Fig. 5.**
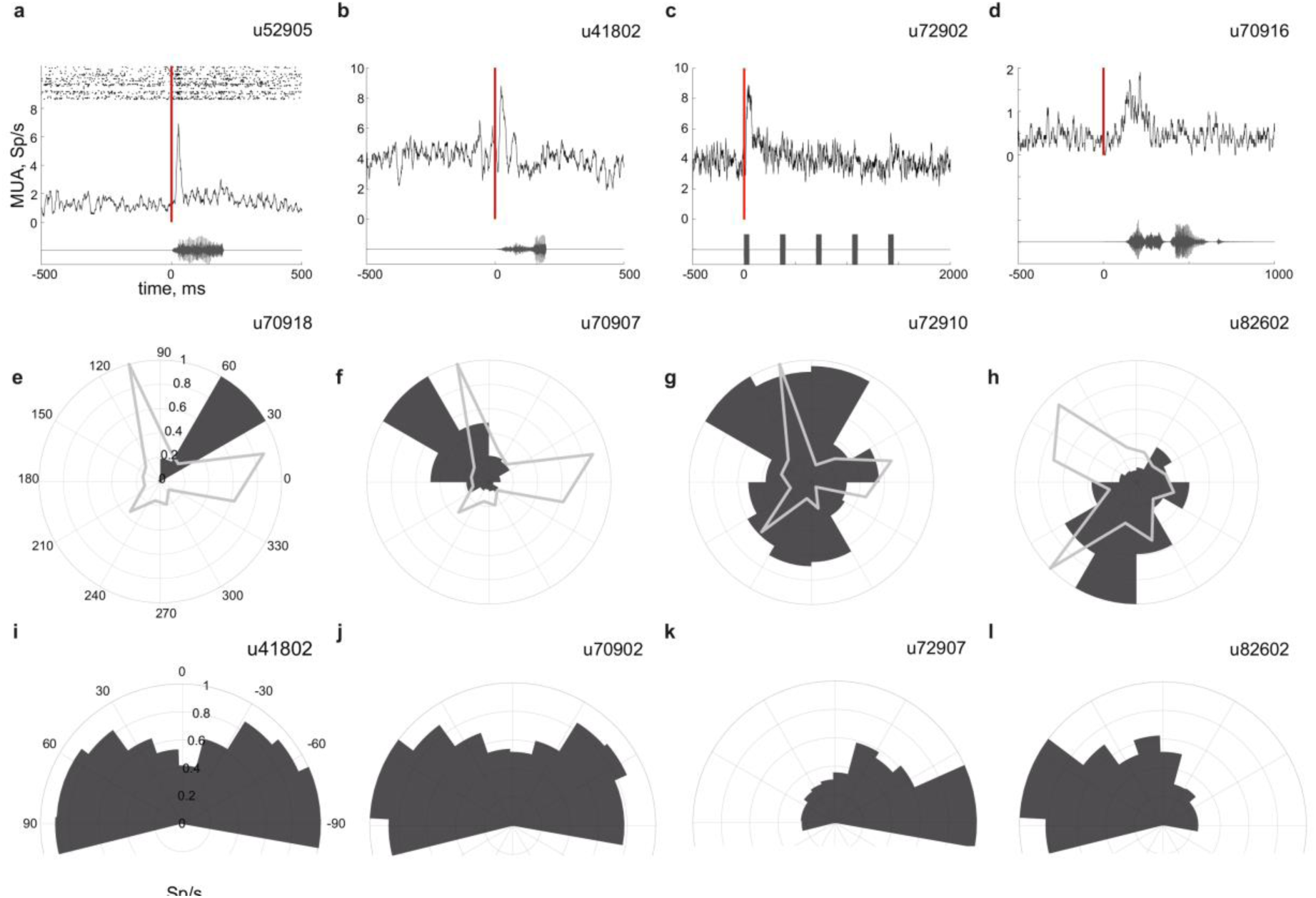
Neural responses during behavior. **(a-d)** Sound driven responses. Mean over one session of the sound driven responses (top) and the oscillograms of the corresponding auditory stimuli (bottom). Red line indicates sound onset. The dots in (a) show the spike times in the individual trials as a raster display. **(e-h)** Location sensitive units. Each panel shows the mean firing rate of a unit while the animal was in one of 12 sectors of the arena (dark gray), and the time spent in each sector (light gray). **(i-l)** Units sensitive to angle between head and body of the animal. The normalized mean firing rate is plotted when the rat exhibited different angles between head and body (dark gray). The last bin on each side includes the data equal and more extreme than the indicated angles. Unit identifiers are marked above each panel.

Remarkably, in both areas, we found units which did not respond to sounds, but whose activity was strongly modulated by other aspects of the task and the state of the rats. Two of these classes of responses are illustrated here.

In Ins, we found units that increased their firing rates selectively at specific locations in the arena (Figs. 5e-h). Since the rats spent most of their time close to the walls of the RIFF in the St+ task, the data are summarized as polar plots. For example, the unit shown in Fig. 5e responded essentially only when the rat was at one specific sector (polar pie chart; 1-way ANOVA, F(1,35060) = 221, p = 7.9*10^-50^). This sector was not preferred by the rat: it spent much more time in the two flanking sectors (line plot in Fig. 5e).

To control for potential confounds, we checked that the firing rate of these units was not determined by the time spent in each sector, and that their location sensitivity was largely independent of their sensitivity to sound, velocity, and head-body angle (Supp. Figs. S5 and S6). Responses in the presence and absence of auditory stimulation showed no significant difference (paired t-test, n.s. for each unit separately, see Supp. Fig. S5). To check the independence of location sensitivity from velocity (or head-body angle), we computed the matrix of mean firing rates as a joint function of spatial position and velocity (or head-body angle). We then computed a rank-1 approximation of this matrix using nonnegative matrix factorization. A good rank-1 approximation to the observed firing rates implies that the two parameters cause largely independent changes in the firing rate of the unit. We then verified that this rank-1 approximation explained an appreciable amount of the structure of the data (see Methods for details). In general, the rank-1 approximation to the observed firing rate accounted for a similar amount of variability as expected from surrogate data, verifying that for the units presented here, location sensitivity was indeed independent of velocity and of head-body angle.

The second type of units illustrated here (Figs. 5i-l) showed sensitivity to the angle between the head and the body of the rat, extracted from the video stream by the data analysis pipeline (see Methods). Such units were found both in AC and in Ins. To control for confounds, we checked that their sensitivity to the relative head direction was not a consequence of their sound responses (paired t-test between angle-dependent mean firing rate in the presence and absence of sound, n.s. for each unit separately, Supp. Fig. S7). The head-body angle sensitivity was independent of location in the RIFF, and of the absolute angles of the rat body and head in space (see Supp. Fig. S8 for full results). The unit presented in Fig. 5h is the same as that presented in Fig. 5l. It showed sensitivity to both rat location and head-body angle, but those sensitivities were independent of each other.

## Discussion

The RIFF is a fully automatic, high-throughput, high-resolution environment for studying the neural basis of behavior in complex tasks and to shape behavior without manual intervention. It collects detailed online behavioral information together with synchronized neuronal recordings. We demonstrated the tracking of behavior and electrophysiology in a precise manner in time and space. Animals learned successful behavioral strategies very quickly. This is somewhat surprising, as learning complex tasks sometimes takes weeks or months (Gronskaya & Behrens, 2019; Miller et al., 2017; Poddar et al., 2013). Potentially, the large environment and the multiple possible actions available at each moment encouraged more extensive exploration (de Hoz & Nelken, 2014; Rosenberg et al., 2021), leading to faster learning.

Virtual reality setups have been developed to flexibly study neural activity during behavior (Aghajan et al., 2015; Dombeck et al., 2007, 2010; Go et al., 2021; Harvey et al., 2009; Hölscher et al., 2005; Keller et al., 2012; Minderer et al., 2016; Radvansky & Dombeck, 2018; Sofroniew et al., 2014). Such systems allow for the stable acquisition of high-quality data in head-fixed animals (Minderer et al., 2016), and can integrate imaging techniques that are harder to apply in freely moving animals (Bjerre & Palmer, 2020). Still, virtual reality is an approximation, may provoke unnatural movement patterns (Huang et al., 2020; Whishaw et al., 2017), and evoke neural responses that differ from those observed in real environments (Aghajan et al., 2015; Ravassard et al., 2013).

Many high-throughput systems for studying freely moving animals have been developed to explore learning and plasticity (Keating et al., 2013), navigation (Fry et al., 2000; Go et al., 2021; Khan et al., 2005; Lim & Celikel, 2019; Tsoar et al., 2011) exploration (Ballesta et al., 2014), social interactions (Ballesta et al., 2014; de Chaumont et al., 2012, 2019; Hong et al., 2015; Matsumoto et al., 2013; Pérez-Escudero et al., 2014; Romero-Ferrero et al., 2019; Shemesh et al., 2013; Straw et al., 2011; Weissbrod et al., 2013) and other behaviors (de Chaumont et al., 2012; Matsumoto et al., 2013; The International Brain Laboratory et al., 2021), and some can track many individuals simultaneously. However, they are often hard to combine with neuronal recordings. Only a few behavioral environments include wireless recording of neural activity from single freely moving animals (Finkelstein et al., 2015; Nourizonoz et al., 2020; Yartsev & Ulanovsky, 2013). The novelty of the RIFF in this emerging field lies in its integrated approach allowing a rich repertoire of behavioral tasks combined with wireless electrophysiology.

The RIFF was designed for maximal flexibility in the investigated behaviors. The ‘tight loop’ of the control program allows for using information from many sensors to react to animal actions at very low latencies. In the St+ task rats learned to move to reward locations before reward availability was signaled, and used stable individual strategies across days. These findings were possible only because of the high temporal resolution and the high-throughput design of the RIFF. This level of detail in the analysis of behavior may be crucial for the analysis of the links between behavior and neuronal activity.

Indeed, we identified unexpected connections between behavior and neural activity. We present examples of neurons in primary and high-order auditory areas that show sensitivity to non-auditory features of the tasks. The highly detailed data collected from each animal allowed us to control these findings for a large number of confounds, such as simultaneously presented sounds, head and body directions, and animal velocity.

In particular, to the best of our knowledge, we provide here for the first time in Ins evidence for non-auditory neurons whose responses depended on the location of the animal in space. Ins receives direct projections from the primary division of the auditory thalamus, the ventral medial geniculate body (Takemoto et al., 2014), and has neurons with frequency-tuned, short-latency responses (Kimura et al., 2010), but is nevertheless spatially separate from the primary auditory cortex (Rodgers et al., 2008).The location-sensitive neurons described here are therefore intermixed with neurons with response properties similar to neurons in the primary auditory cortex. The evidence provided here adds to other recent studies that described location-sensitive neurons in many structures outside the hippocampal system (Jankowski et al., 2015; Jankowski & O’Mara, 2015), including visual (Long et al., 2021), somatosensory(Long & Zhang, 2021), retrosplenial (Mao et al., 2017), and premotor (Yin et al., 2018) cortices.

Importantly, the RIFF represents a concept that can be extended by adding more sensors, for example a motion capture setup (Gardner et al., 2021; Marshall et al., 2021), more actuators for interacting with the animals, and electrodes with hundreds of contacts (Steinmetz et al., 2021). The RIFF shifts the emphasis from single, reduced measures of behavior such as success and failure, into high rate, detailed information streams that are a better fit to the neural dynamics of the mammalian brain.

## Materials and methods

### Behavioral setup

The arena, 160 cm in diameter, is composed of 18 modular sections (28 cm width, 50 cm height - aluminum skeleton and 5 mm thick black matte Perspex sheets; one section is illustrated in Fig. 2b). The floor is made of two sheets, tightly connected by screws. The arena is mounted on a 5 cm high supporting structure, of the same size and shape of the floor sheets with depressions for the connection between the two sheets as well as for the insertion of gridded floors (for possible foot shock punishment). The arena is electrically grounded with a cable running around the outer perimeter near its bottom.

Each of the 18 sections contains three panels where additional equipment can be inserted. In the RIFF, the outer panels of every third section contain two nose-poke ports (Fig 2; a total of 12 ports in 6 sections, SPECIAL.090-SE v1.0, LaFayette-Campden Instruments, Loughborough, UK). One of the two ports in each section is connected to a liquid pump and the other to a pellet dispenser. In addition, all 12 ports are connected to an airpuff valve (80210raas, LaFayette-Campden Instruments, Loughborough, UK). At the top of each panel with a port, a free field speaker (MF1, Tucker Davis Technologies, Alachua, FL, USA) is mounted in a custom-made holder with an adjustable angle. Such a section, with its two ports and two loudspeakers, is termed an ‘Interaction area’ (IA) in the paper.

The ports are controlled by a combination of commercial software (ABET II, LaFayette-Campden Instruments, Loughborough, UK) and a custom-written MATLAB program (The MathWorks, Inc., Natick, MA, USA). Signals to and from the ports interface with a digital I/O card (NI PCIe-6509 96 ch, National instruments, Austin, TX, USA) through specially designed hardware. The digital I/O signals are monitored and modified by the custom-written MATLAB program that implements the MDP.

### Auditory stimulation

In both tasks described here (St+ task and L/D task, described below), a small set of sounds was used. These sounds were synthesized ahead of time and stored in computer files. Pure tone stimuli were synthesized in MATLAB. Word-like stimuli were created using the following procedure. Words were selected from free online recordings (http://shtooka.net). They were then processed using the STRAIGHT Vocoder (Kawahara et al., 2008) and MATLAB. For sounds in the St+ task the spectrotemporal envelope was extracted by STRAIGHT and shifted above 1 kHz. For the sounds in the L/D task, the spectrotemporal envelope extracted by STRAIGHT (Kawahara et al., 2008) was shifted and stretched along the frequency scale, so that the envelope levels at frequencies 0.1 kHz and 20 kHz of the original sound were shifted to frequencies 1 kHz and 40 kHz (with linear interpolation on a logarithmic scale in between). The resulting spectrotemporal envelope of each sound was used to resynthesize it using the original pitch contour.

During the experiment, all sounds were loaded to memory. When a sound had to be presented, it was transduced to voltage signals by a sound card (M-16 and MADIface USB, RME Audio Interfaces, Munich, Germany), attenuated (PA5, Tucker Davis Technologies, Alachua, FL, USA), and then played through one or more of the 12 speakers, driven by stereo power amplifiers (SA1, TDT) .

The acoustics of the arena were studied in detail (Kazakov et al., 2018). In short, the arena produces two major reflections, one from the floor and the other from the wall behind the animal. The dominant reflection from the floor creates spectral modulation with a period of 5 kHz and depth of about 5 dB. Absolute sound levels at 0 dB attenuation were about 100 dB SPL at 2 kHz, going down to about 85 dB SPL at 20 kHz. Sound levels were rather stable as a function of location in the arena (varying by <5 dB between the center of the arena to the walls) and of loudspeaker (standard deviations across loudspeakers at the center <5 dB; at the wall <8 dB).

No correction was applied for this variation.

### Animals

The experiments were carried out in accordance with the regulations of the ethics committee of The Hebrew University of Jerusalem. The Hebrew University of Jerusalem is an Association for Assessment and Accreditation of Laboratory Animal Care (AAALAC) accredited institution. Seven adult female Sabra and Sprague Dawley rats (N = 5 and N = 2, respectively, weight: at least 200 gr) were used for the experiments described here (Envigo LTD, Israel). All efforts were taken to create a low-stress, rat-friendly living environment enabling experimental animals to freely express their innate behaviors. Upon arrival, animals were housed in groups (2-3) or individually (depending on experiment) in the same SPF room in which the RIFF is situated and where all experiments took place. The temperature (22 ± 1 °C) and humidity (50 ± 20%) were controlled and room was maintained on a 12-h light/dark cycle (lights on from 07:00 to 19:00 h)

### Behavioral training

For the first 7-10 days rats were habituated to the experimenter. During this period, rats were handled frequently and introduced to new types of foods and flavored water which were later used as rewards in the experimental arena. The procedure was aimed to decrease stress and prevent neophobia towards rewards given during behavioral training. Rewards consisted of palatable 45 mg food tablets in six flavors: basic company flavor, banana, bacon, chocolate, peanut butter and fruit punch (diet AIN-76A: RodTab45MG; RodTabBan45MG; RodTabBcn45MG; RodTabChoc45MG; RodTabPbtr45MG; RodTabFrtP45MG5TUL, TestDiet, Richmond, IN, USA). Rats received also 4 types of fluids: mineral water, 4% sucrose in mineral water, 0.1% saccharine in mineral water, and 1:1 mixture of 4% sucrose and 0.1 % saccharine solutions (sucrose and saccharin, Sigma-Aldrich, St. Louis, MO, USA).

In order to motivate animals to work for food, rats were food-restricted up to 85% of their ad libitum body weight. Rats were subjected also to mild water restriction before behavioral sessions (4-12h). Experiments were carried out 5 days a week. During the weekend animals had free access to standard rodent food and unsweetened mineral water.

#### Multiple strategy task (St+)

In this task, rats had to move to specific locations at defined times indicated by distinct sound sequences, in order to obtain rewards. The experiment consisted of a repeated iteration over three events: Attention (ATT) - Target (TGT) - Feedback (FDB). Each event was associated with a specific sound that was played whenever the event occurred. The exact paradigm is described in Fig. S1a. The ATT event consisted of the presentation of an ATT sound - a 50 ms broadband 2 KHz periodic sound that was repeated 5 times at intervals of 0.3 s. It was played from all 12 speakers simultaneously. The position of the rat was determined 2.5 s after the ATT event, and one of the IAs was selected as the target location. The selection was done according to the current and previous position of the rat (see below for details). Then a TGT sound was presented from the target location. Each IA was associated with a specific TGT sound. The TGT sounds consisted of a sequence of six 50 ms long narrowband sounds with inter-sound intervals of 250 ms covering a range of 1-16 kHz. Six different permutations of those sequences were assigned as target sounds to each IA for the whole experiment (target area 1: 1, 1.7, 3, 5.2, 9.2, 16 kHz; target area 2: 1.7, 16, 1, 9.2, 5.2, 3 kHz; target area 3: 3, 1.7, 1, 16, 9.2, 5.2 kHz; target area 4: 5.2, 3, 1.7, 1, 16, 9.2 kHz; target area 5: 9.2, 16, 5.2, 1, 3, 1.7 kHz; target area 6: 16, 9.2, 5.2, 3, 1.7, 1 kHz). After the TGT event, the rat had to nose-poke into one of the two ports at the target IA. Nose-poking within 20 seconds after the TGT event was rewarded by food pellets or water, depending on which of the two ports of the IA were poked. Reward was provided only if the first nose-poke was in the correct IA, and only to that first nose-poke. The first nose-poke (in a correct or wrong location) elicited the FDB event at the target IA, consisting of a sound indicating whether the location was correct or not (word-like stimuli derived from the word “correct” or “mistake”, respectively). A timeout (20 seconds without poking after a TGT event) was also followed by an error FDB sound. After each FDB event, an inter-trial period of 3 seconds led to the next ATT event.

The target locations were selected in the following way. The arena was subdivided (Fig. S1a) into A zones, in the immediate vicinity of the IAs; B zones, adjacent to the A zones but 17 cm towards the center; C zones, separating the A/B zones of nearby IAs; and a large D zone in the center of the arena. The A and B zones were slightly wider (33 degrees) than C zones (27 degrees). When the rat was in the D zone, nothing happened. If the rat was in one of the A/B/C zones, an ATT sound was presented (3 s following the last FDB event). The rat had to make a choice where to go within the next 2.5 s. ‘A’ choices consisted of the rat remaining in the same A zone in which it received the last reward, in which case the target was presented from the next A zone in the clockwise direction, or by a move to another A zone, which was then selected as the target. In these cases, the rat got 1 reward (pellet or liquid). ‘B’ choices consisted of a move to a B zone, in which case the next target consisted of the adjacent IA and the reward was 3 times larger than when making ‘A’ choices. The rat could therefore cycle between a B zone and the adjacent IA ports, harvesting rewards from the same port. Finally, the rat could move to a C zone, in which case a random target was selected and the reward was randomly chosen at 1-4 times the reward of the ‘A’ choices. In the data shown here, rats learned an ‘A’ strategy - moving around the arena from one IA to its neighbor, circling the arena. In fact, the rats rapidly learned the timing structure of the task, and often moved to the next A zone before the ATT sound was presented (Fig. 4d).

The rats were trained in three phases. During phase 1 rats were trained to poke for reward. Multiple food and water restricted rats were placed together in the arena for about 12 hours for five consecutive days. The sounds were not presented at this point. The rewards were delivered in a fixed ratio (FR) schedule. The number of pokes needed to get reward was increased every day as follows: FR2, FR7, FR10, FR12, and FR15.

In phase 2, rats established instrumental and Pavlovian associations with relevant sounds and their sequences. Rats were trained individually, each during a 70 min long session performed once a day. The training was carried out for 30 days. The sound level was increased gradually to avoid suppression of behavioral responses and to introduce animals gradually to the relatively high SPL of the sounds during the main experiment. Each day, only 1 IA was accessible for a rat during the session. Interactive ports in all other 5 areas were closed by custom blinds. Consecutive sessions were carried out in areas 1 to 6 and such a cycle was repeated 5 times. There were 3 types of sounds presented in this phase. The rat started the trial by poking into the accessible port. Each poke (photo beam break) evoked the 50 ms broadband 2 KHz periodic sound. After five successful pokes with sound, the sixth poke triggered reward delivery (1 unit) and immediately the target sound (TGT) was played from the two speakers above the accessible IA. The TGT sound was followed by 1 s of silence and the word-like stimulus derived from the word “correct” played from two speakers above the accessible IA. An obligatory 2 s interval followed before the RIFF reacted again to nose-pokes of the rat. Thus, during that stage, while sounds were associated with reward, it was the nose-poke that triggered the sounds rather than the reverse (as in the main task).

Phase 3 was the main experiment as described above. Rats performed a 70 min long experimental session once every day.

The behavioral data in Fig. 4 was collected during the first two days of Phase 3 of the task. However, the electrophysiological data shown in Fig. 5 was collected after one additional refinement of the task (phase 4). In randomly selected 10% of the trials, a warning sound was presented (the phrase, composed of word-like stimuli: “do not go’’, played from all speakers). The rats had to avoid accessing the ports in order to avoid air-puff, starting 1.5 s after the warning sound (to let the rat retract its nose from the port within in case it was poking when the warning sound was presented). An airpuff was triggered by a nose poke for pokes that occurred during the next 3.5 s. If the rat successfully avoided poking any of the ports, a safety sound (the phrase, composed of word-like stimuli: “it’s safe”) was presented from all speakers. Otherwise, an airpuff was delivered and the phrase “you failed” (composed of word-like stimuli) was presented from all speakers. After safety or punishment sounds, a new trial was initiated with ATT sound.

#### Localization/discrimination task (L/D)

In this task, each of the IAs was associated with a sound. The sounds were 200 ms excerpts of 6 different words (the word “here” in 6 languages: English, French, German, Italian, Polish and Russian) in a female voice, processed as in the Auditory Stimulation section. The rats had to go to a central circular area to initiate a new trial (Fig. S1b). When a crossing into the central area was detected, one of the 6 IAs was randomly selected by the program and its associated sound was played once every 2 seconds from both speakers for up to 10 times (20 s in total), or until the rat poked in any port. A poke in one of the two ports of the IA from which the sound was played resulted in a reward (food or water, according to the poked port). A wrong poke or a timeout (20 s without a poke) resulted in the termination of the trial.

The rats were habituated to the RIFF during 2 nights (24 hours in total). During habituation, The RIFF was divided into 6 equal sectors centered on the IAs (Supp. Fig. S1b). Rats were placed in the RIFF and were able to explore freely. Whenever the rat entered an active sector, the associated sound was played from the speakers of the IA in the sector. The sound repeated every 2 s until the rat poked in one of the ports of the IA and got a reward, with a timeout after 2 minutes. After a nose poke, the current sector and its two neighbors became inactive and the three other sectors became active, so that the rat had to cross to one of the three sectors on the opposite side of the RIFF in order to initiate a new interaction.

Following the habituation, the rats were exposed to increasingly stricter versions of the main task in 12 hour overnight daily sessions. For the first 6 training sessions the rats were allowed to poke in more than one port before trial termination (10 pokes in the first and second sessions, decreased to 6 and 4 for one session each, 2 pokes for 2 sessions, then down to 1 for the rest of the experiment). In addition, the requirements for initiating a trial became stricter: on the first training session, the central area had a radius of 50 cm, on the 2nd session of 40 cm, and from the 3rd session on the central area had a radius of 30 cm. Before electrode implantation rats were switched from 12 hour overnight sessions to 3 hour morning sessions.

### Electrodes

Rats were chronically implanted with 32-channel silicon probes (ASSY-116_E-2, Cambridge Neurotech, UK). Prior to implantation thin flexible ground wire was soldered to the electrode’s PCB ground contacts. The electrodes were aligned and glued to the microdrive shuttle with epoxy (Nano Drive, Cambridge Neurotech, UK). The microdrive with the electrodes and connector were held by a custom holder attached later to a stereotaxic apparatus for implantation. The moveable parts of the microdrive were covered with paraffin oil to prevent possible leak of dental cement between them. Before implantation, the electrodes were cleaned by washing them with a 4% tergazyme solution in purified DDW water (Alconox, Jersey City, NJ, USA) and afterwards carefully washed in purified DDW to remove all residues of tergazyme. Before implantation the electrodes and the holder were disinfected in a UV sterilizer.

### Surgery

The implantation of the silicon probes was performed in two stages: (1) preparation of the base for the implant and (2) implantation of microelectrodes into the brain tissue.

#### Preparation of the base

Rats were initially anesthetized in an induction chamber with sevoflurane (8% in oxygen, Piramal Critical Care Inc., Bethlehem, PA, USA). The head was shaved and they were placed in a stereotaxic instrument with a mask for gas anesthesia (David Kopf Instruments, CA, USA). Sevoflurane concentration was slowly adjusted to a level of 2–2.5% and maintained at this level throughout the surgery. A surgical level of anesthesia was verified by the lack of a pedal-withdrawal reflex and slow, regular breathing rate. Body temperature was controlled by a closed loop heating system with a rectal probe (Homeothermic Monitoring System, Harvard Apparatus, MA, USA). The eyes were protected with sterile eye drops for dry eyes (Viscotears Liquid Gel, Carbomer: polyacrylic acid 2 mg/g, Berlin, Germany) and the skin on the head was disinfected with a povidone-iodine solution (10%, equivalent to 1% iodine, Rekah Pharm. Ind. Ltd., Holon, Israel). To prevent postoperative pain, rats received during the surgery subcutaneous injection of Carprofen 50 mg/ml (5% W/V) in a dose of about 12 mg/kg (Norocarp, Norbrook Laboratories Limited, Newry, Co. Down, Northern Ireland).

A 1.5–2 cm longitudinal cut of the skin on the head was made and the dorsal surface of the skull was exposed. The opened skin was stretched and the eyes were closed. The left temporal muscle was pulled away to expose also the lateral surface of parietal and temporal bones. The connective tissue covering the bones was removed and bones were treated with a 15% hydrogen peroxide solution (Sigma Aldrich Inc., St. Louis, MO, USA) which was washed off with sterile saline after approximately 10-20 s. When the surface of the skull was clean and dry, a reference point for the entry point of the recording electrodes was marked (Insular cortex (InsC): AP = -1.0 mm, ML = -6.1 mm; Primary auditory cortex (A1): AP = -5.1 mm, ML = guided by landmarks on the lateral surface of parietal and temporal bones). Subsequently, 7 small holes for supporting screws were drilled and screws were tightly screwed into the frontal, parietal and interparietal bones. Ground wire, soldered previously to one of the screws, was placed in the left frontal bone. The screws were fixed together and to the bone first with resin and then acrylic dental cements (Super-bond C&B, Sun Medical, Moriyama, Shiga, Japan; Coral-fix, Tel Aviv, Israel) forming a base of the implant. At the selected electrode implantation site, a thin polyimide tube was placed on the skull vertically and cemented with the rest of the implant base. The tube served as a guide to the implantation site for the next surgery. The free end of the ground wire was twisted and covered with a polyethylene cap cemented to the rest of the implant.

The wounds were cleaned and treated in situ with antibiotic ointment (synthomycine, chloramphenicol 3%, Rekah Pharm. Ind. Ltd., Holon, Israel). The skin was sutured in the anterior part of the implant with one or two sutures (Nylon, Assut sutures, Corgémont, Switzerland) to stretch the skin around the base of the implant. The skin around the wound was cleaned and covered with a povidone-iodine solution (10%). The rats received intraperitoneal injection of the antibiotic enrofloxacin 50mg/ml (5% W/V) in a dose of 15 mg/kg diluted with saline to 1 ml (Baytril, Bayer Animal Health GmbH, Leverkusen, Germany). After surgery, animals were housed individually to prevent them from chewing the implants. Carprofen or other similar NSAID dissolved in palatable wet food was provided at the home cage for the first few days after surgery. The rats were allowed at least 1 week of recovery post-surgery before restarting behavioral training.

#### Implantation of silicon probes

When the wounds were completely healed following the first surgery (14-28 days), the recording electrodes were implanted. To minimize tissue damage, a small craniotomy was made, drilling solely through the base of the implant, and thus leaving the healed skin intact to accelerate recovery and reduce the pain.

As previously, rats were initially anesthetized in an induction chamber with sevoflurane (8% in oxygen). After induction, rats were transferred to a stereotaxic instrument with a mask for gas anesthesia. Sevoflurane concentration was slowly adjusted to the level of 2–2.5% and maintained at this level throughout the procedure. The eyes were protected with sterile eye drops for dry eyes (Viscotears Liquid Gel, Carbomer: polyacrylic acid 2 mg/g, Berlin, Germany) and body temperature was controlled by a closed loop heating system with a rectal probe.

The dental cement above the implantation site marked by polyimide tube was removed gradually using a dental drill until the skull was exposed. The craniotomy was performed by drilling, and a 0.4-0.8 mm long slit in the dura was gently resected. The electrodes were slowly inserted into the brain tissue using a single axis micromanipulator (MO-10, Narishige, Tokyo, Japan). The craniotomy was sealed with paraffin oil and elastic silicone polymer (Duragel, Cambridge Neurotech, UK). The microdrive and connector were fixed to the base of the implant with acrylic dental cement. A ground wire was soldered between the base and the electrodes connector and covered with acrylic dental cement. To mechanically stabilize the connection between the implant and the recording device, an additional supporting connector was cemented to the implant (6 pins of 853 Interconnect Socket, MILL-MAX MFG. CORP., New York, USA). A custom plastic enclosure with a screw cap was cemented for implant protection. The robust design of the implant protected the microdrive and the electrodes against mechanical stress while the rats were moving, improving recording stability (Supp. Fig. S9d-f). The weight of the whole construct didn’t exceed 11.5 g.

At the end of the surgery, to prevent postoperative pain, rats received a subcutaneous injection of Carprofen 50mg/ml (12 mg/kg). Rats received intraperitoneal injection of enrofloxacin 50mg/ml (15 mg/kg) diluted with saline to 1 ml. Rats were allowed at least 3 days of recovery post-implantation before recordings.

### Wireless electrophysiology

Two wireless recording systems were used: (1) modular 64 channel neural logger (RatLog-64, Deuteron Technologies, Jerusalem, Israel) and (2) 64-channel wireless transmission system (TBSI W64, Triangle BioSystems International, Durham, NC, USA).

#### Neural logger

For the experiments described here we used a single processor board and a single amplifier board to record from 32-channel silicon probes at a sampling rate of 32 kHz. The data were saved on a 64 GB micro SD card, and copied to a computer after each session. The logger was equipped with an audio microphone and 9-axis motion sensor. An electrically insulated, compact enclosure for the logger, with a separate compartment for a battery, protected the logger from mechanical shocks (Supp. Fig. S9). The enclosure included an interconnector, to protect the board connector against mechanical damage caused by attaching and releasing over hundreds of recording sessions (Supp. Fig. S9a). The interconnector was also used to rotate the device about 34 degrees backwards from the vertical axis, to allow the rats free movement and free access to the ports in the IAs (Supp. Fig. S9b). The enclosure of the logger had a supporting connector (6 pins of 852 + 853 Interconnect Socket, MILL-MAX MFG. CORP., New York, USA) that matched the supporting connector attached to the implant, in addition to a 36-pin Omnetics connector (A79029-001, Omnetics Connector Corporation, Minneapolis, MN, USA) for the electrical signals from the electrodes. The total weight of all elements was approximately 16 g. The load was balanced, to avoid pulling the rat head in any direction. Before every recording session, the device was additionally secured with autoclave sticky tape (Sigma-Aldrich, St. Louis, MO, USA) to prevent loosening of the connectors during the 2-3 hours duration of the recording sessions. Maximal recording time using a 300 mAh battery was about 3 hours (LiPo battery 582030).

#### Wireless analog transmission system

The system consisted of an analog transmitter and receiver (TBSI W64, Triangle BioSystems, Durham, NC, USA). The output signal of the receiver was routed to a data acquisition system (AlphaLab SnR, Alpha Omega) and digitized at a sampling rate of 44 kHz. The recordings were made from 32-channel silicon probes as above. A small interconnector with a battery holder was prepared in the lab. The headstage was placed horizontally with a battery holder on its left side enabling animals to freely access to all locations in the experimental arena, and in particular to the IAs. The total weight of all elements was approximately 13.5 grams of well-balanced load. Before each recording session the device was additionally secured with autoclave sticky tape (Sigma-Aldrich, St. Louis, MO, USA) to prevent loosening of connectors contact during long active behavioral sessions (up to 12 hours). Maximal recording time using a 260 mAh battery was about 11.5 hours (LiPo battery 601240).

### Recordings

Neural signals were recorded in reference to a ground placed in the frontal bone. For the logger, the analog bandpass filter was set to 10-7000 Hz or 300-7000 Hz depending on the experiment. For the wireless system, the low frequency of the bandpass filter was set to 0.07 Hz. Recording sessions took place five days a week and implants were checked daily. Spiking activity was screened immediately after each recording session was finished. The electrodes were kept in the same position as long as spiking activity was detected on many contacts. When signals deteriorated, the animals were briefly sedated with sevoflurane and the electrodes were lowered in steps 25, 50 or 100 µm into the brain tissue. Electrodes were moved typically every 1-7 days. Electrodes were never moved up.

### The video imaging system

A custom video system tracked the trajectory of the animal in real-time at 30 frames/second (Supp. Movie S3). The localization latency was less than 30 ms and a compact storage format resulted in data volume of 4.3 GB/hour. The system robustly tracked rats of different sizes and was unaffected by the presence of a head implant.

#### Hardware

A monochrome camera (DMK 33G445 GigE, TheImagingSource, Bremen, Germany) was mounted on the ceiling above the center of the arena. A wide-field lens (T3Z3510CS, Computar, North Carolina, US) covered the whole arena. A LED ring (50 cm in diameter, CN_R64, NanGuang, Chenghai, China) placed around the camera provided a uniform diffuse illumination (Fig. S10a). The luminance of the ring was set to the minimal level that was sufficient for exposure time of 25 ms.

The camera was connected to a computer running the Windows operating system with a GigE cable that supplied power and transmitted the grayscale 1280x960 pixel images. The triggering pulses were sent from the camera by a Hirose cable to the digital input section of a multifunction device (RX8, Tucker-Davis Technologies). The RX8 sub-sampled the incoming trigger train from 30 Hz to a 1 Hz pulse train and output it to the main recording unit (AlphaLab SnR, AlphaOmega). These 1 Hz triggers also powered a LED that was placed outside the arena but inside the field of view of the camera (Supp. Fig. S10b). The LED state was used to synchronize the images with the rest of the recorded data during the experiment.

#### The real-time module – image acquisition and rat localization

A custom graphical user interface (programmed in MATLAB) was used to launch the tracking program and monitor its activity in real time (Supp. Fig. S10c). A single background image of the empty arena was acquired during the initialization of the tracking program. The real-time tracking routine executed an acquisition-localization tight loop: Every time a new image was acquired from the camera, the rat was located by subtracting an image of the empty arena from the current image of the arena with the rat inside. The difference image highlighted the area where the rat was located. The location of the rat was estimated by fitting an ellipse to the rat pixels and computing its center of mass. The x and y coordinates of the estimated rat center of mass were sent to the computer that controlled the experiment over an Ethernet connection and are also stored as metadata.

To reduce data storage, the video stream was saved in a compact form. Each rat image was cropped around the rat center of mass, since only this area was expected to change from one frame to the next. The original image could be approximately reconstructed by placing the cropped rat image on top of the background image at the location indicated by the coordinates of the center of the rat, stored in the metadata file. In addition, one of the corners of the bounding box was replaced by a small rectangle that showed the LED (Supp. Fig. S10d). The images and the metadata were stored in a compact database at the rate of 4.3 GB/h. The processing of a single video frame required less than 30 ms.

### Post-experiment data processing

After the conclusion of each experimental session, data were processed by a custom, automated pipeline written in MATLAB (tested with version 2020b, code publicly available under github.com/jniediek/RIFF_publication/tree/main/processing). Data processing started with the extraction, alignment, and merging of the time stamps from all data acquisition devices. Time stamps were generated for changes in internal states, the acquisition of video images, behavioral events, sound presentations, and neural recordings. The digital lines of an AlphaLab SnR system (Alpha Omega) were used as the main synchronization hub for aligning the triggers coming from all devices. Each device that generated timing triggers had at least one of those triggers also channeled in parallel to a digital line on the main synchronization hub, and the timing events that were co-registered on both devices were used to align all other events from that device on a common timeline. Thus, the output files of the SnR system were used as the main timing files during post experimental data processing. All behavioral events (nose-pokes, food/water delivery, airpuffs) as well as the video images were parsed, and the extracted information stored in MATLAB files. Finally, the neural recordings were processed, detecting and sorting spikes (see next section).

### Spike Sorting

We developed a pipeline that processes an experimental session of 3 hours in about 10 minutes, based on the Kilosort2 spike sorting program (https://github.com/MouseLand/Kilosort2). The processing was predominantly autonomous, requiring only a few minutes of manual curation for every session.

Spike sorting began by removing common modes from the neural recordings - the mean activity of all spatially proximal recording channels, such as all channels on a single shank of a silicon probe (Supp. Fig. S3a.1-2). Large artifacts were identified by computing the standard deviation over electrodes at each point in time, and locating peaks in the resulting time series. Such artifacts were mostly generated by rat movements. Signal sections around each of these peaks were zeroed by multiplication with a 10 ms window with 2 ms rising and falling edges (Supp. Fig. S3a.3). The total recording duration that was zeroed during a typical experimental session was about 5%.

Spikes were detected and clustered by Kilosort2. In order to process large amounts of data, we developed a fast and almost automatic pipeline, which can be complemented by detailed manual curation where necessary. Kilosort2 was used to produce the initial set of candidate spike clusters. Automatic rejection of noise clusters was performed according to two criteria (Supp. Fig. S3b): The dominant channel of the cluster template has exactly one minimum and one maximum in its voltage trace, and on each electrode the voltage at the template endpoints had to be close to 0 (in consequence, confining expected spike duration to <1.8 ms). For each unit that passed this test, an inter-spike-interval histogram was computed alongside the amplitude and firing rate histograms, heat-map of raw spike waveforms, and smoothed time series of spike amplitude and firing rate (Supp. Fig. S3c). All those were printed to an image file that summarizes the quality of that unit.

Lastly, a manual labeling interface allowed the experimenter to sort the units into well isolated ones, multi-unit clusters or noise. This interface displayed the images with the quality evaluation information and stored the user-inserted labels with the data. A typical recording session of 1-2 hours with 32 electrodes produced typically a few tens of non-noise units, so that the manual labeling could be completed in several minutes.

### Feature extraction from rat images – convolutional neural network

Pose information was extracted from the saved video stream, offline, in two steps: First, the nose, the base of the neck and the base of the tail were located on the image by a convolutional neural network (Supp. Fig. S2a, Supp. Movie S1). Then, the relevant angles were computed by geometric calculations (Supp. Movie S2, green arrow represents the body direction). The neural network was trained de-novo, including data labeling, model training and time optimizations of the inference process.

#### Network architecture

The input to the model is a square grayscale image of the rat, as produced by the rat tracking module. The output consists of three pairs of Cartesian coordinates, representing the three points of nose, neck and the base of the tail on the rat image. The model is thus a multivariate regressor that predicts 6 continuous variables.

We used a custom convolutional neural network architecture, following the recent advances in data-driven computer vision (Supp. Fig. S3b). Six convolutional layers are used, each followed by a batch-normalization layer and a rectifier non-linear activation function. A max-pool layer operates after every second convolutional layer. The last layer is a linear readout from the previous activation map stack. The network has 0.3 million trainable parameters.

#### Database creation

We used a custom programmed graphical user interface (written in MATLAB) to manually mark the locations of the nose, neck and base of the tail on each image of the training set. The final training database included 1500 labeled images of 4 rats of different ages, with and without a head implant.

#### Network training and inference enhancements

The training database was augmented to produce additional training examples and to emphasize the irrelevant degrees of freedom in the image space. The augmentation included rotation of images by multiples of 90 degrees, horizontal mirror flips, adding small amounts of noise to pixels or labels, and rigid translations of images by a few pixels along the horizontal and vertical axes. The labels were re-adjusted for each augmentation type to correctly represent the updated body part location. This augmentation procedure increased the original dataset by a factor of 64, from 1500 to 96,000 training examples.

The network was initialized with the default Kaiming distribution (He et al., 2015), trained with the Adam optimizer (Kingma & Ba, 2017) on 256 images in each batch. The input images were subsampled to 50x50 pixels. The network was trained for 2 hours on a NVIDIA GTX 1080Ti GPU (NVIDIA, California, US).

After the training was completed, the network could be used to predict the labels of previously unseen images. Prediction used image augmentations that resemble those used during the training: An input image was processed 4 times, rotated by 0, 90, 180 or 270 degrees. The predicted 4 labels were corrected for the corresponding image rotation and averaged to obtain the final label for the original image. The high temporal correlations of the image stream were leveraged to increase the accuracy by smoothing the labels with a uniform window of 7 frames, removing large, biologically implausible jumps.

An hour of a typical experiment produced ∼105 images, which were labeled by the trained model in ∼20 seconds.

#### Computation of allo- and ego- centric directions

Once the locations of the nose, neck and the base of the tail are predicted by the network, they are used to calculate the direction of various body parts. The main body direction is defined as the vector from the base of the tail to the base of the neck. The head direction is defined as the vector from neck to the nose point. The head turn angle is calculated as the difference between the head direction and the body direction, with 0 corresponding to the three points being along a single line. The range of head turn angles was typically between -30 and 30 degrees.

### Final database formats

The data processing pipeline parsed each experiment into a set of data files, metadata descriptors and statistical visualizations. This format is uniform and allows an abstraction of the experiment parameters such as the type of neural recording system, the stimulus types, and so on. All behavioral and neural events are stored in a lossless manner, allowing maximal temporal resolution. The storage footprint of this format is about 12GB/hour.

For rapid analysis and visualization, a temporally coarse-grained version is generated by discretizing the timeline into a 10 Hz grid, and aggregating the separate data files into a single data table (stored in one file). This format uses about 1% of the storage of the raw storage (<0.1GB/hour). Each data row in the resulting data table has a time-stamp that indicates its start time on the time-axis of the original data. These time-stamps can be used as indexing variables for tracing the data back to the high-resolution files.

### The experiment visualizer

All processed data from an experiment can be loaded into an interactive user interface that presents the recorded experiment as an interactive movie (at a rate of 10 Hz, see Fig. 3f). The raw neural data is loaded along with the detailed time information of the sorted spikes, making it possible to view both the firing rate of each unit and the times of individual spikes with a 1 ms resolution.

The visualizer was used for multiple purposes. During the development of the experiment, replay of the experiment with 100 ms resolution made it possible to verify that the experimental logic was correctly implemented and executed by the hardware, and the rat indeed performed the required sequence of actions. The visualizer was also used for verifying the collected data; the temporal resolution of the visualizer makes it possible to detect potential data collection malfunctions such as synchronization errors. During data analysis, the visualizer was used as a tool for understanding fine details of rat behavior and for observing the progress of the learning process. Lastly, the visualizer can plot the raw neural traces at any time point of the experiment, making it possible to observe the relationships between spiking activity and other events in the RIFF. Figures 3c and 3f show images produced by the visualizer. The visualizer was used to generate Supp. Movies S4 and S5.

### Statistical analysis

Exploratory analysis of the dependence of spike trains on any of the measured parameters was conducted using linear mixed effects models (MATLAB function fitlme). Explanatory variables included location and kinematic parameters, body and head direction parameters, and sounds presentations. The models were fitted to the firing rates of the individual units collected in all experimental sessions of each rat. To check the effect of any of these parameters on the neuronal responses, it was used as a fixed effect (in order to account for a non-zero mean across the neuronal population) as well as a random factor depending on each recorded unit.

These random factors are highly regularized (they are estimated under the assumption that they are instances of a Gaussian variable with a data-dependent covariance matrix (Verbeke & Molenberghs, 2000)), and are therefore conservative estimates of the dependence of the firing rate of each specific unit on the parameter of interest.

We selected for further study units with prominent random effects. Prominent random effects were defined as effects whose magnitude was greater than the standard deviation of the corresponding fixed effect. The logic behind this choice is based on the observation that for a unit that has a prominent random effect, the dependence of its firing rate on the parameter of interest is different from that of the population mean. This method identified sound-sensitive neurons, but in addition it identified rat location, velocity, and head-body angle as variables that affected the activity of many units in all rats.

We highlight in this paper specifically location sensitive units and head-body angle sensitive units. While units could show sensitivity to multiple variables, we verified the independence of the effects of the variable of interest from all other variables. Thus, all examples shown here were not the result of a spurious correlation due to another primary dependence. We illustrate such tests with an example of location sensitivity. For each unit, the average firing rate as a joint function of location and one additional variable (for example, velocity) was computed, usually using a 10-by-10 grid of bins, and represented as a matrix. In order to check the independence of the tuning to the two variables, a rank-1 approximation was computed for this matrix using nonnegative matrix factorization (MATLAB function nnmf, see Supp. Fig. S6). We then compared the matrix containing the average firing rates with its rank-1 approximation. To measure this, we divided the L2 norm of the residual by that of the original matrix. Small numbers represented good approximations and therefore a high level of independence. To follow standard usage, we subtracted this number from 1 and used that as the analog of the fraction of data variability explained by the rank-1 approximation.

Since the original matrix was estimated from a finite sample, it is expected to have a high rank even if its ideal structure has rank 1. Therefore, two methods were used to estimate the expected fraction of data variability explained by a rank-1 approximation to a noisy data matrix whose underlying structure is also of rank 1. In the first method the original data was used for bootstrapping: the firing rates observed within each location bin were resampled, and assigned to random velocity bins according to their observed probability. The mean for each one of the matrix entries was calculated. The resulting 10-by-10 matrix was processed in the same way as the original data. This process was repeated for 100 times, the fraction of explained variability averaged and compared to that of the original matrix. In the second method, the rank-1 approximation was used to generate surrogate data with Poisson distribution, using the same number of counts in each bin of the original matrix. This matrix was processed in the same way as the original data, and the fraction of explained data variability from this process was compared to that of the original data.

All the units were also tested for the effects of stimulus driven responses (Supp. Figs S5 and S7). The mean responses with and without sound presentations were computed as a function of the relevant parameter (in each location bin for the location-sensitive units, for each head-body angle bin for the head-body sensitive units). The mean firing rates in the presence and absence of sound were then compared using a paired t-test.

In all statistical analyses, significance was defined as p< 0.05. Exact data are reported within the results section, and include the model, the statistic tested and the p-value.

### Data and code availability

The data described in this manuscript are available at github.com/jniediek/RIFF_publication, along with the MATLAB code to create the corresponding figures, to process raw data recorded in the RIFF, and to operate the RIFF during experiments. A sample session recorded from the RIFF is available at figshare, https://doi.org/10.6084/m9.figshare.15082971.

## Acknowledgments

We thank Tamir Scherf for his help with the design and implementation of the image processing neural network, Dr. Tamar Ravins and the staff of the Authority for Biological and Biomedical Models at Hebrew University for their excellent assistance in assuring high standards of animal welfare. This work was supported by AdERC grant GA-340063 (project RATLAND), by F.I.R.S.T. grant no. 1075/2013 from the Israel Science Foundation. JN was supported by a DFG Research Fellowship (ref. NI 2012/1-1; project number 442068558). IN holds the Milton and Brindell Gottlieb Chair in Brain Sciences.

## Author contributions

IN conceived and supervised the project. MMJ and AP designed and constructed the RIFF, designed the behavioral paradigms with a contribution of JN, and collected data. AP wrote the software for implementing Markov Decision Processes in the RIFF. MMJ designed and developed the chronic neural implants. AK developed the rat tracking, spike-sorting, image processing modules and the experiment visualization GUI. JN, AK and AP developed the analysis pipeline for the data from the RIFF. MMJ, AP, AK and JN jointly brought the system to life. AP and JN analyzed the behavioral and neural data presented in this paper.

## Supplementary Materials

**Fig. S1.**
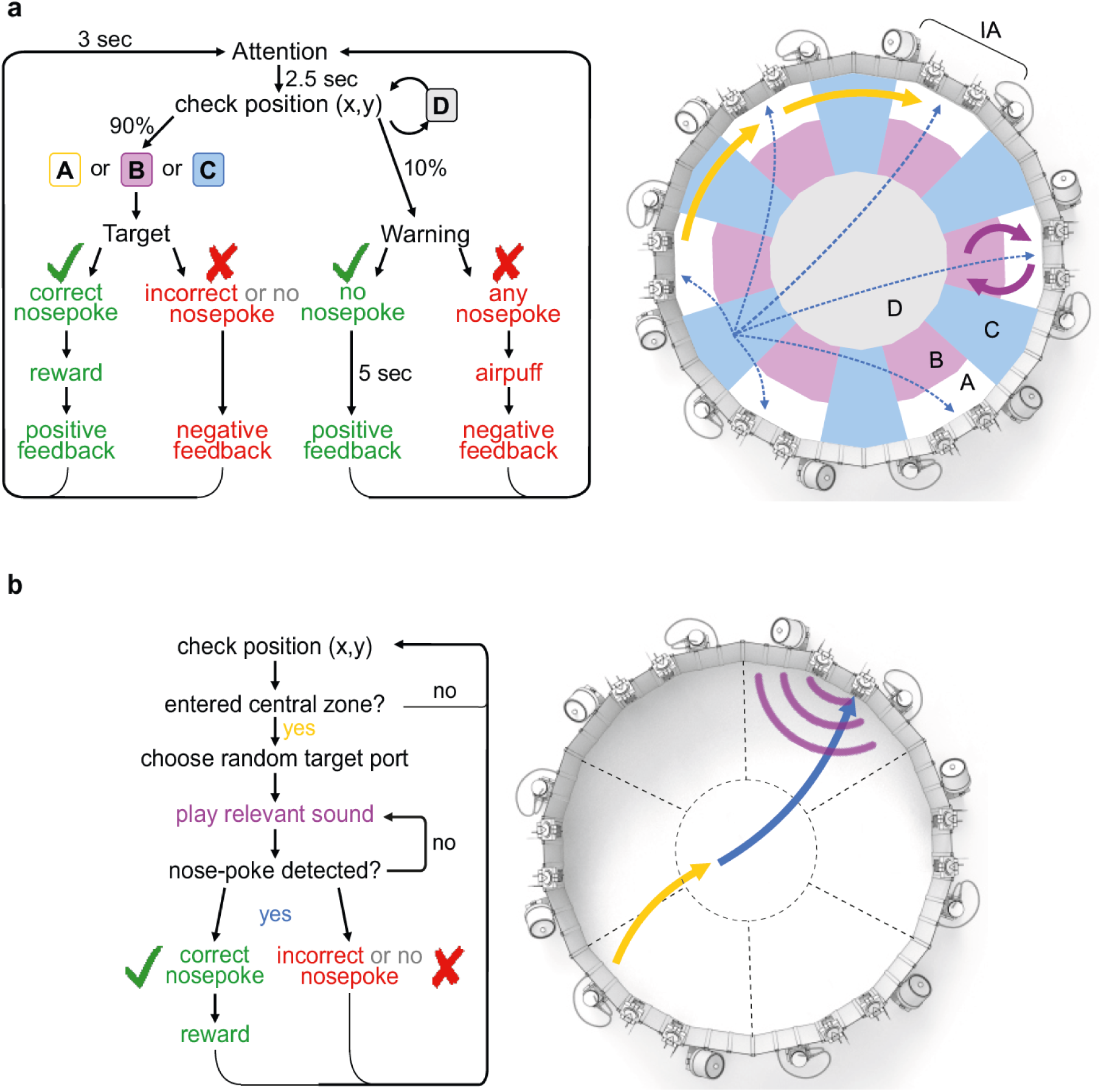
Structure of the two tasks described in the paper. **(a)** The multiple strategies task (St+). On the left, a flowchart of the experiment is displayed as a real-time loop. At the check position stage, three strategies were available to the rat, marked as A, B and C. These strategies are illustrated in the diagram of the arena on the right, using the same color code. The A strategy consisted of moving from one interaction area to another (usually a neighboring area), which was then selected as the next target. The B strategy consisted of cycling from an A area to the associated B area, which led to the selection of the same A area as the next target. The C strategy consisted of moving to a C area, in which case a random port was selected as the next target. **(b)** Diagram of the localization/discrimination task (L/D). Flowchart of the experiment real-time loop (left) and the corresponding events in a diagram of the arena (right), plotted with the same color code.

**Fig. S2.**
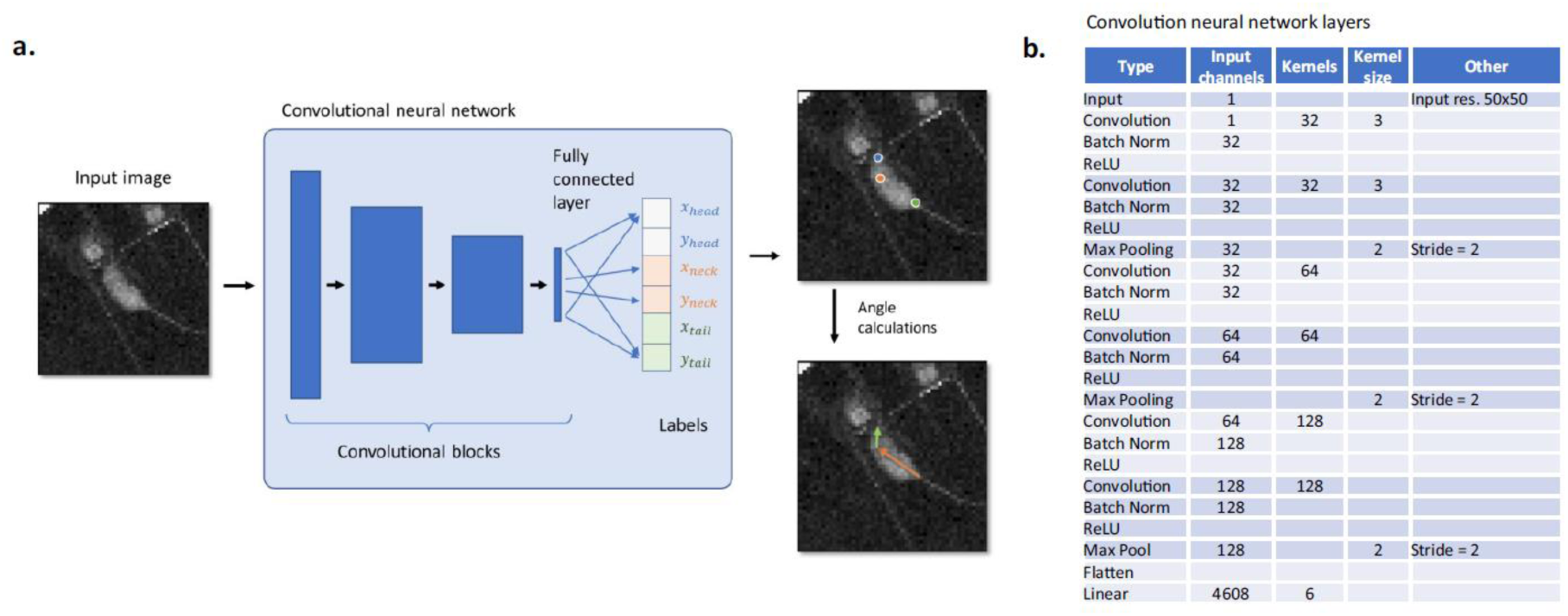
Feature extraction from the video images. **(a)** A feed-forward convolutional neural network estimates the locations of the head, the base of the neck and the base of the tail for each input image. These three markers are then used for the calculation of the body and head angles (bottom right image). **(b)** The table details the custom architecture of the neural network, which is optimized for the grayscale rectangular input, reducing the number of parameters of the trained model and decreasing the inference times.

**Fig. S3.**
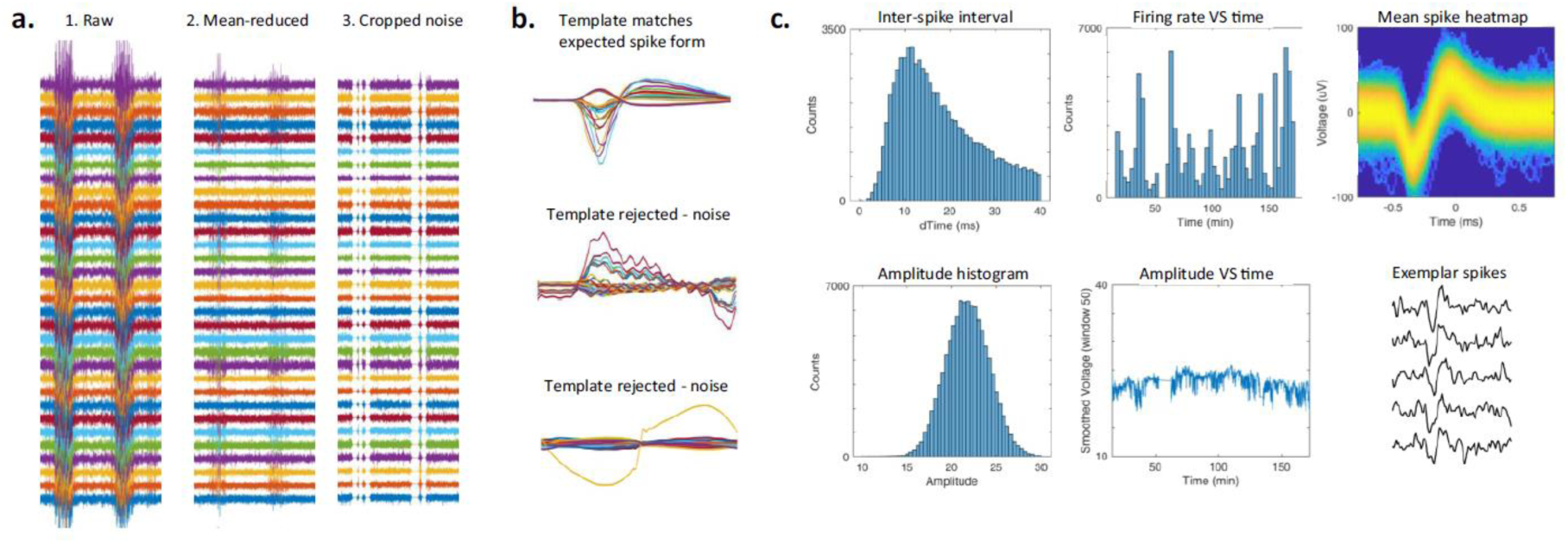
Neural data processing and spike sorting. **(a)** Extracellular neural recordings in freely behaving rats (32 simultaneously recorded channels) include periods of high noise (left panel). Noise components that are common to all channels can be largely removed by subtracting the average waveform (middle panel). The remaining noise segments are identified by their amplitude and by their high variance across channels, and are zeroed (right panel). The resulting neural data is then processed by Kilosort2. **(b)** Implausible spike shapes in the clusters detected by Kilosort2 are automatically detected. The top example was automatically identified as a spike, while the bottom two examples were identified as noise. **(c)** Statistics of the neural activity are produced for each cluster and used for manual classification.

**Fig. S4.**
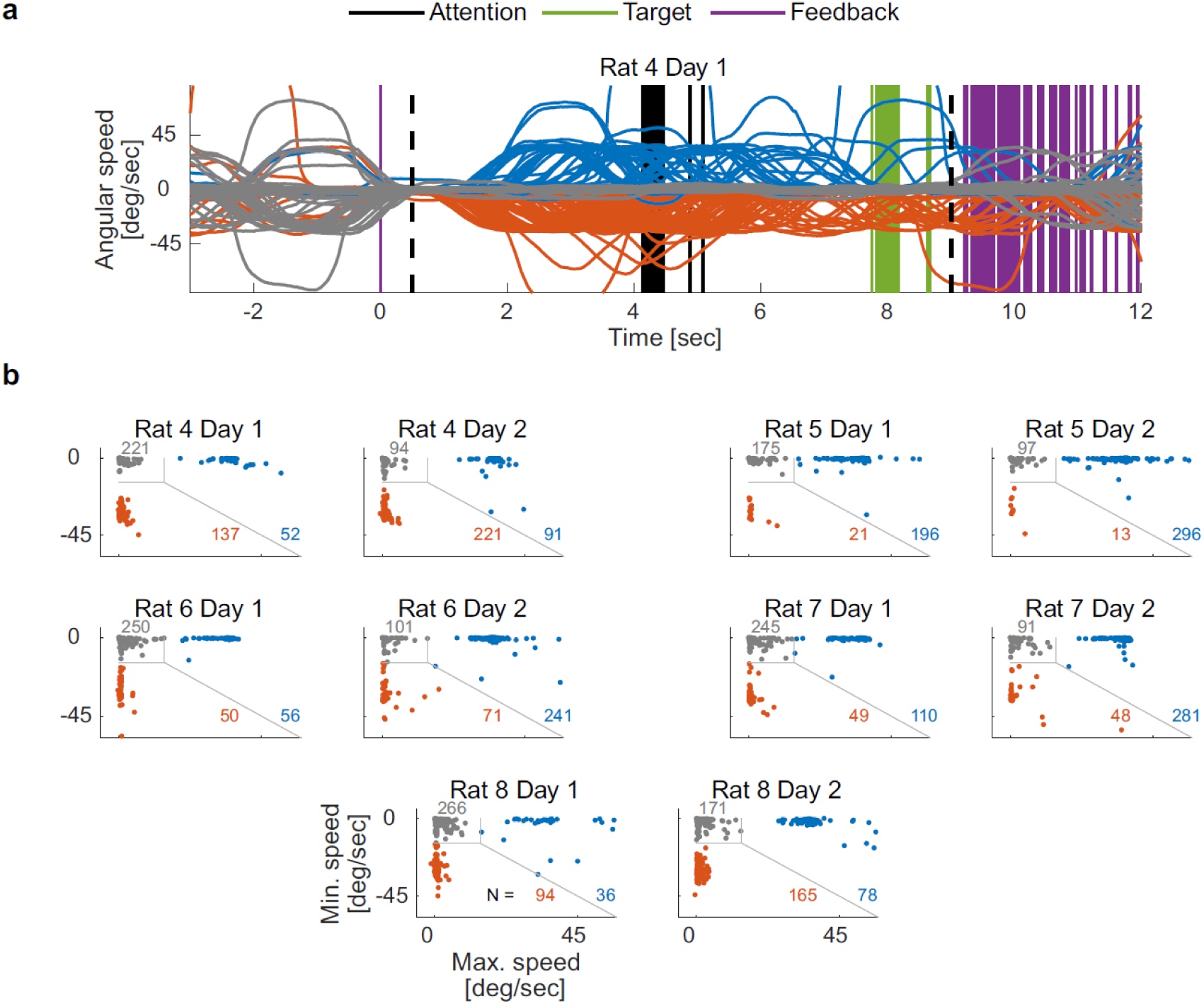
Classification of trials into three types. **(a)** Angular running speed for all trials performed by rat 4 on day 1. For each trial, the maximal angular speed in the clockwise and counterclockwise direction was extracted in a time window lasting from 0.5 seconds to 9 seconds following the feedback sound of the previous trial. A trial was classified as “Sit” (gray lines) if the absolute value of the angular speed never exceeded 0.25 radians/s (14.3 degrees/s), otherwise as “Run clockwise” (blue lines) or “Run counterclockwise” (red lines), according to the direction with the higher maximal speed. **(b)** Trial clusters were clearly separated in all rats. Scatter plots show each trial of each rat and each day according to the maximal angular speed in the clockwise and counterclockwise directions. Colors as in (a). The number of trials of each trial cluster are indicated.

**Fig. S5.**
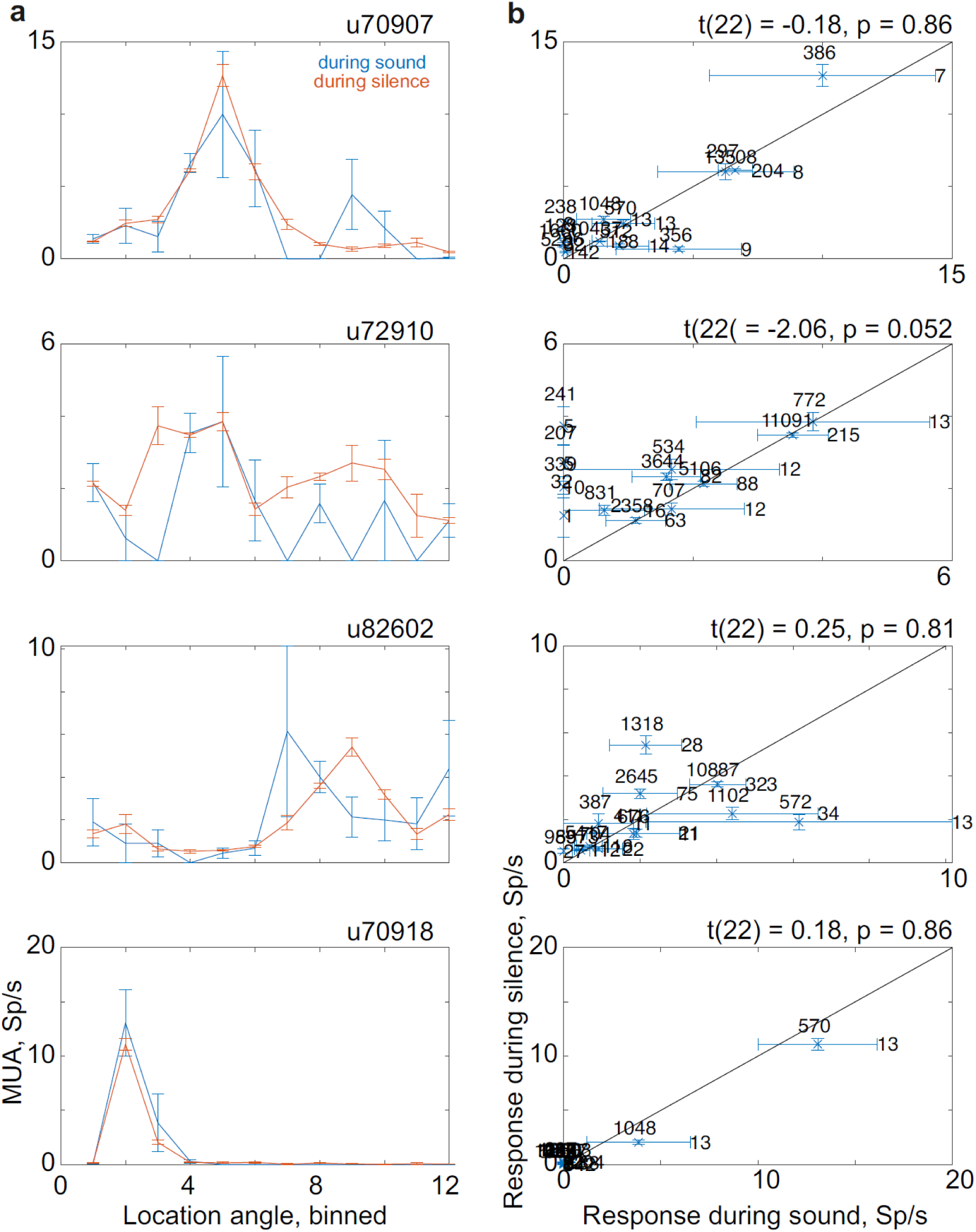
Joint sensitivity of the units in Figs. 5e-h to location and sound. **(a)** Mean responses in the presence (blue) and absence (red) of sound. Error bars are s.e.m. **(b)** Scatter plots of the data as in (a). Horizontal error bar indicates s.e.m during sound presentation, vertical error bar indicates s.e.m during silence. The number of instances from which the mean and s.e.m were derived are indicated on the right of each point (during sound presentation), and on top (during silence). In case no sound presentations occurred in a location bin, the data is plotted on the y axis.

**Fig. S6.**
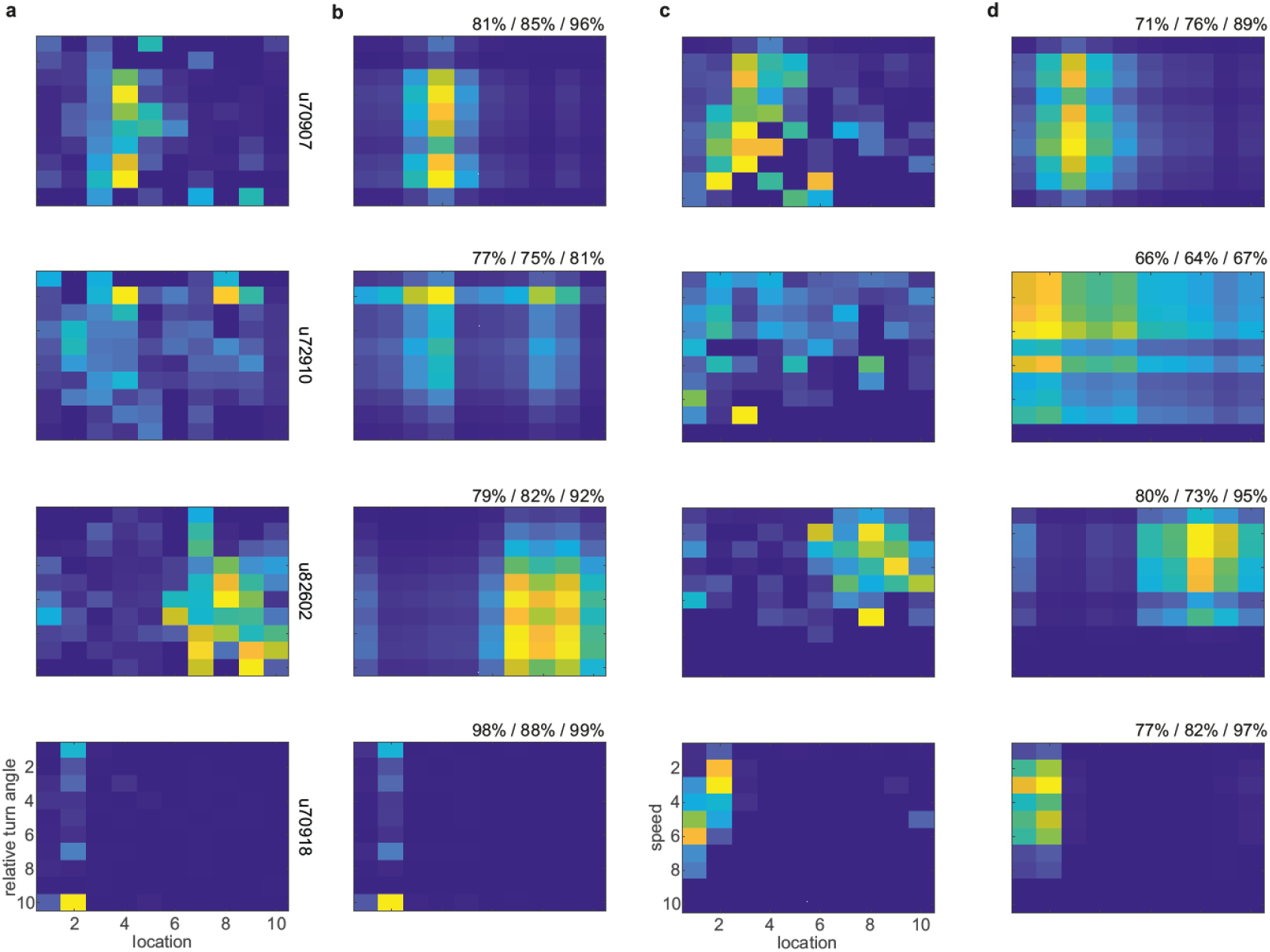
Joint sensitivity of the units in Figs. 5e-h to location and head-body angle or velocity. **(a)** Mean firing rates for location (abscissa) and relative head-body angle (ordinate). **(b)** Non-negative rank-1 matrix approximation of the matrices in (a). Top right corner: the fraction of the data variability explained by the rank 1 approximation for the original data / bootstrapping method / Poisson distribution approximation (see Methods for details). **(c)** Mean firing rates for location (abscissa) and velocity (ordinate). **(d)** Same as (b) for the matrices in (c).

**Fig. S7.**
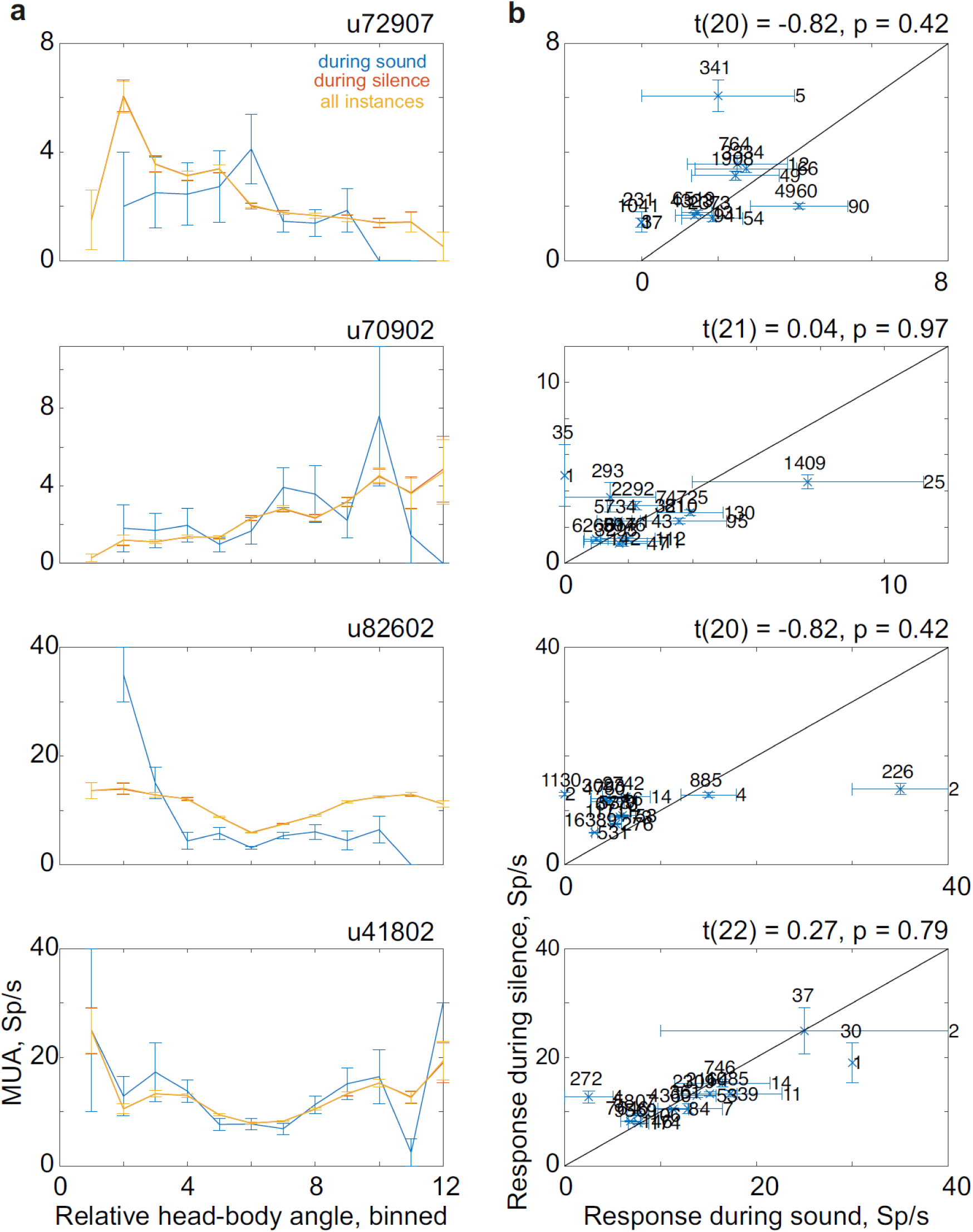
Joint sensitivity of the units in Figs. 5i-l to location and sound. Same format as Fig. S5.

**Fig. S8.**
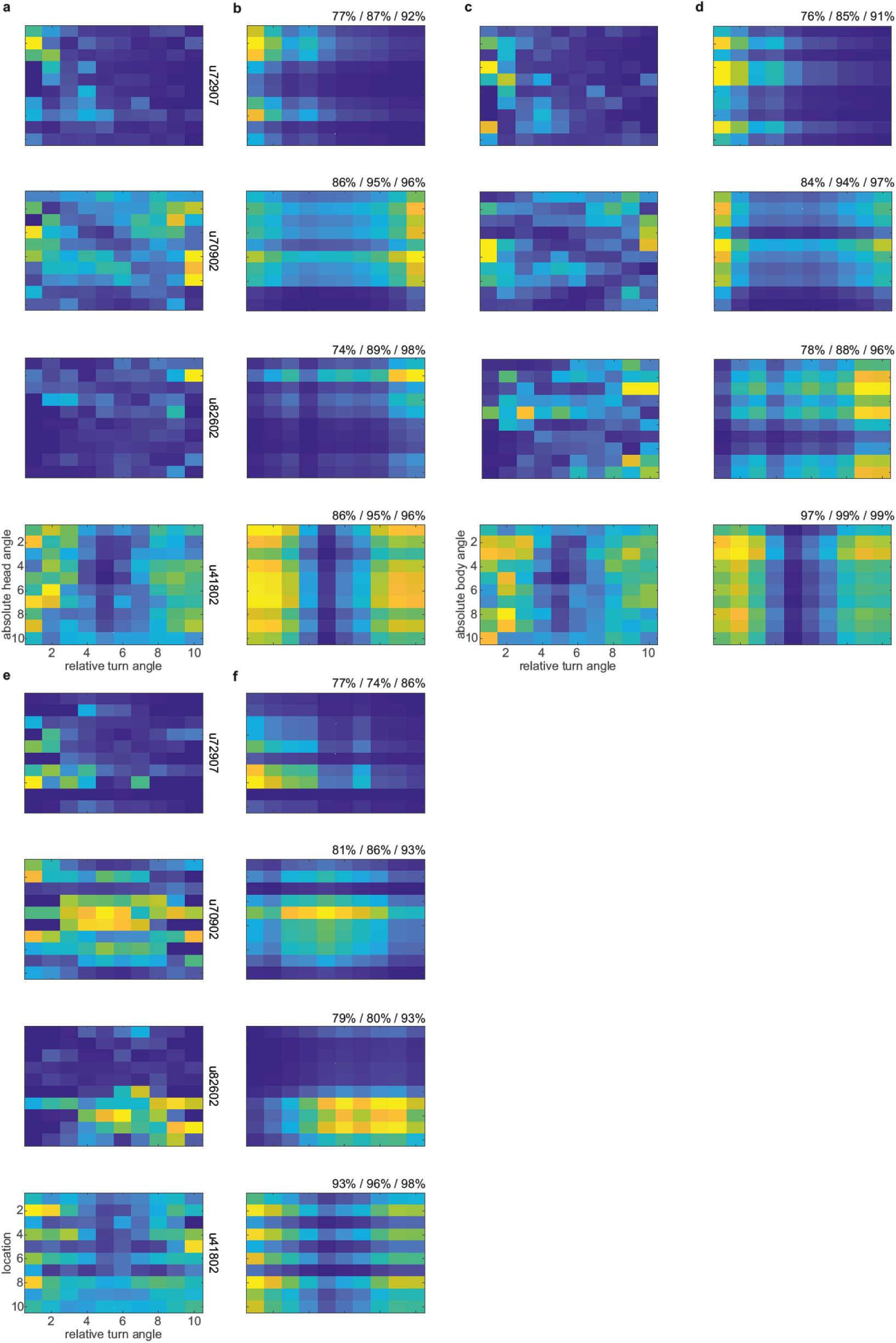
Joint sensitivity of the units in Figs. 5i-l to head-body angle and absolute head angle, absolute body angle, or location. Same representations as in Fig. S6. The abscissa represents head-body angles, while the ordinate represents absolute head angle ((a) and (b)), absolute body angle ((c) and (d)), and location ((e) and (f)).

**Fig. S9.**
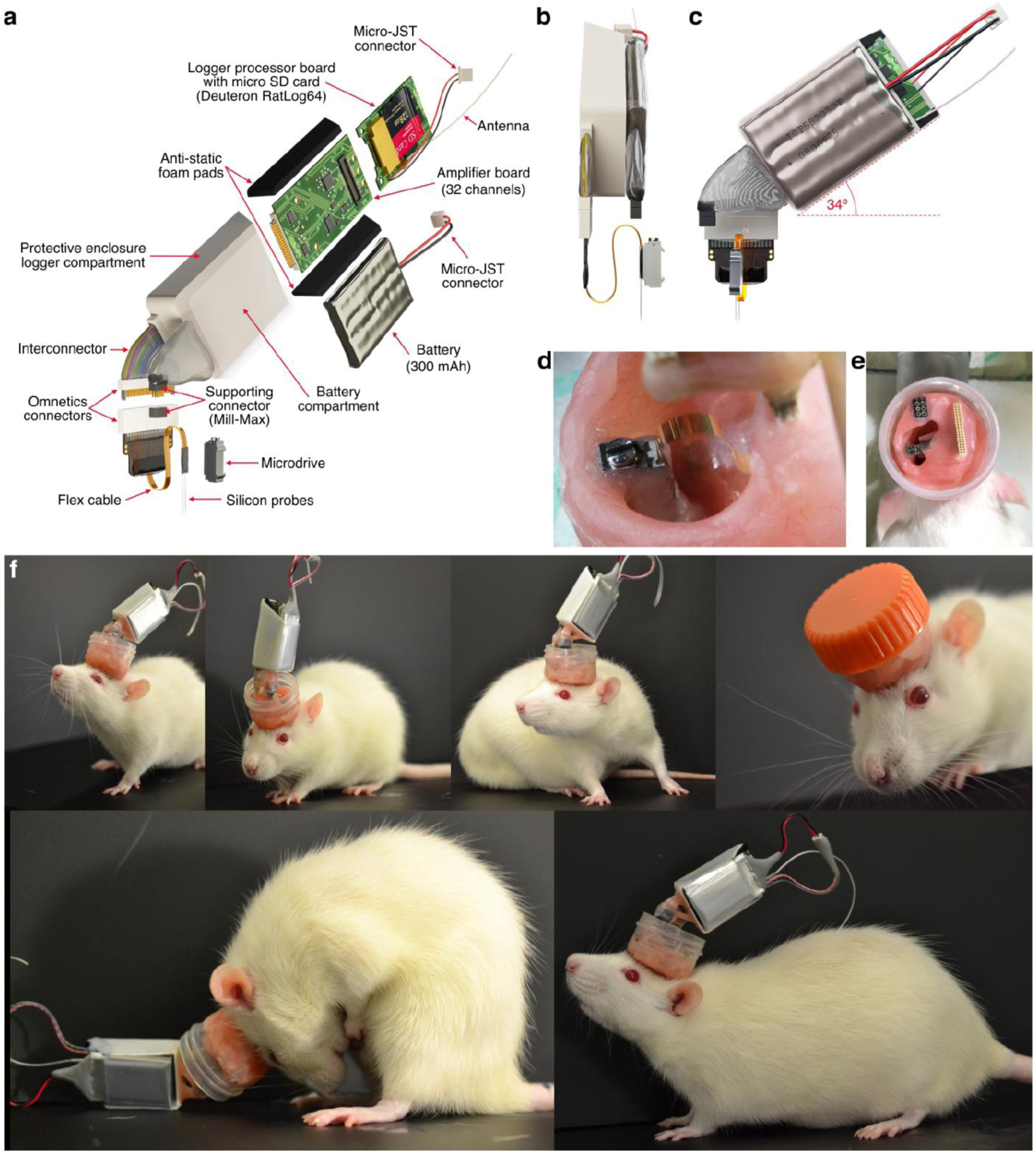
An approach for chronic wireless recordings in rats. **(a)** The neural logger and battery are in the protective plastic case that can be attached to the electrode’s connector. The 300 mAh battery is placed in a separate compartment, connects to the logger through a micro- JST connector, and can be easily changed during the experiment. Anti-static foam pads are placed on the sides of the logger components (amplifier and processor boards) to protect the logger against mechanical shocks. The protective case has a 36-pin omnetics connector matching that on the 32-channel silicon probe, as well as a small Mill-Max connector which mechanically stabilizes the case during recording sessions. The silicon probe is mounted on the Microdrive. **(b and c)** Side and front views of the recording set. The device is inclined to the back in order to allow rats natural movements and undisturbed access to ports. **(d)** The silicon probe is mounted on the Microdrive cemented to the skull. The moveable parts of the implant are covered with paraffin oil. The flex cable of the probe is bent to provide a long travel distance for the electrodes. **(e)** Finished implant with protective enclosure. **(f)** Female rats with a 32- channel moveable silicon probe implant and the wireless data logger in the case with a battery. The recording set enables natural movements, is easily carried by the rats, and is well protected against mechanical shocks. The enclosure can be closed with a plastic cap (orange) to protect the implant in the home cage.

**Fig. S10.**
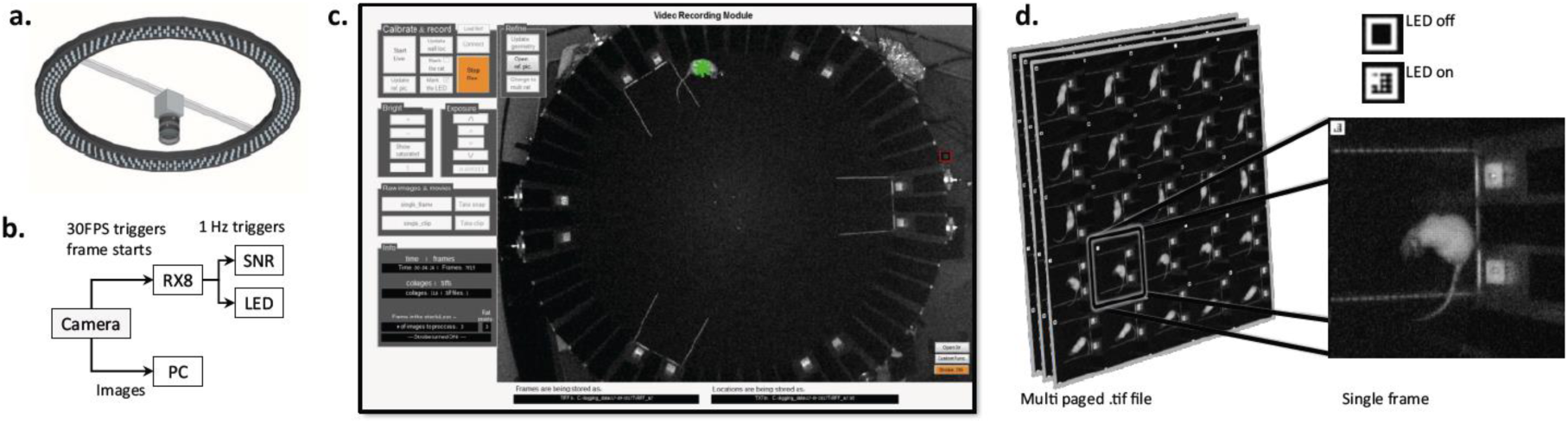
The real-time imaging module. **(a)** Diagram of the camera (DMK 33G445 GigE, TheImagingSource) and the light ring, mounted on the ceiling above the arena’s center. **(b)** Synchronization diagram of the image stream. Triggers that indicate frame acquisition were sent to a digital processor that sub-sampled them from 30 Hz to 1 Hz. The 1 Hz triggers were then simultaneously recorded on the common synchronization hardware, and also powered a LED in the field of view of the camera. **(c)** Graphical user interface of the real-time imaging module. The LED is marked by a small red square on the right side of the arena. The rat center of mass is marked by a green asterisk. **(d)** For efficient data storage, rat images were cropped around its center of mass and stored in a multi-page .tif file. A cropped image of the LED was stored in the upper left corner of each image, allowing for time synchronization during the post-processing steps.

**Movie S1. Automatic detection of 3 rat body points.** An artificial neural network automatically marks the locations of the nose, base of the neck and the tail on a rat image (yellow, red and blue dots, respectively).

**Movie S2. Calculation of the body direction of the rat.** Green arrow indicates the calculated body direction from the base of the tail towards the base of the neck (blue and yellow points from Supp. Movie S1).

**Movie S3. Rat tracking module.** The location of the rat is determined in each video frame, at 30 FPS, as it freely moves inside the arena (green asterisk marks the center of mass of the rat). The location information is transmitted to the main control computer, while the video frames are stored and used to extract additional behavioral features during the post-processing.

**Movie S4. Exemplary behavior of an expert rat in the St+ task at its full complexity.** Rat successfully switches between zones A1 and B6, avoids the air puff in response to the warning sound, and after the safe sound, it continues to use A and B zones while missing an opportunity coming from the C zone (see Supp. Fig. S1a). The St+ task provides substantial freedom to the rats, allowing them to optimize their behavior with respect to their individual preferences for size and type of rewards while effectively avoiding the punishment. Red circles mark active speakers; yellow circle indicates detected nose-poke; green circle indicates dispensed reward. White arrows indicate automatically extracted head and body directions. Dots behind the rat indicate previous location, color indicates speed (blue - slowest, red - fastest). The legend in the top right corner indicates the area where the rat is located. The name of the target sound appears while it plays.

**Movie S5. Exemplary behavior of an expert rat in the L/D task.** Rat goes to the center zone of the arena to initiate a trial with a sound presentation (see Supp. Fig. S1b and Methods for a full description of the task). All markers are the same as in Supp. Movie S4.

